# From genes to collective modes: biological constraints shape metabolic evolution

**DOI:** 10.64898/2026.03.11.711104

**Authors:** Aedan Brown, Sarah Datta, Pankaj Mehta, Brian Cleary

**Author notes:** A.B, S.D, P.M, B.C conceptualized the reasearch and performed preliminary simulations. A.B and S.D performed large scale simulations and analyzed experimental data. A.B, P.M, B.C wrote the manuscript. The authors have no competing interests. This manuscript was deposited on biorxiv under a CC-BY-NC 4.0 International license.

## Abstract

Population genetics has been successful in explaining how drift, selection, and mutation shape allele frequencies. However, we still lack an understanding of how biological and evolutionary constraints arising from genotype-phenotype maps shape evolutionary dynamics. To explore this question, we developed a new framework for simulating the evolution of an inherently polygenic and epistatic trait - metabolism - combining population genetics with flux balance analysis. We found that the evolution of metabolic networks exhibits surprisingly simple, reproducible dynamics. We identify evolutionary collective modes (EvCMs) – linear combinations of genes with high, constant selective pressure – as organizers of the long-term dynamics and the origin of this simplicity. EvCMs arise through the interaction of physical constraints, evolvability, and the requirements for growth. We developed a theoretical framework to predict EvCMs from large-scale metabolic models and found quantitative agreement between theory and simulation of *E. coli* central metabolism. Inspired by these results, we re-analyzed mutational data from Lenski’s long-term evolution experiment and found compelling evidence for the existence of EvCMs. This work suggests that biological constraints encoded in genotype-phenotype maps play an important role in shaping evolutionary dynamics, offering a new perspective on complex trait evolution — one that shifts focus from individual genes to collective modes.

---

Most features of life today were shaped by evolution. Yet, understanding how complex biological traits evolve remains a central challenge in biology. Population genetics excels at mathematically modeling how mutation, selection, recombination, and genetic drift change gene frequencies in populations over time (1–6). However, the abstract nature of population genetic models often obscures the central role played by biological constraints in evolutionary dynamics (4, 7–10). For this reason, understanding the evolution of higher-level traits such as metabolism, which involve complex biochemical networks that are regulated and controlled by large numbers of interacting genes (11–13), requires new theoretical approaches that explicitly incorporate genotype-to-phenotype maps into evolutionary processes.

To investigate evolution in the presence of realistic genotype-to-phenotype maps that are polygenic and epistatic, we performed evolutionary simulations of metabolic networks using a sequential fixation model where mutations arise in a population one at a time, and either fixate or go extinct depending on the fitness advantage they convey to an organism (1, 14, 15). Our simulations show that evolutionary dynamics are strongly shaped by the metabolic and genetic constraints arising from the genotype-to-phenotype map underlying metabolism. The reason for this is that the growth rate of an organism, and hence the fitness advantage a mutation conveys, is a highly non-local, collective property. Physical conservation laws, such as mass conservation (*i*.*e*., flux balance (9)), place strong constraints on the types of mutations that can fix, constraining evolutionary trajectories. A surprising feature of our simulations is that despite this complexity, the evolutionary dynamics of metabolic networks are predictable and can be understood in terms of “evolutionary collective modes” (EvCMs) – linear combination of genes that have high constant selective pressure. The EvCMs organize the long-term evolutionary dynamics and encode the direction of evolution in genetic space. The emergence of EvCMs is especially noteworthy because the selective pressure on individual genes, including those that constitute the EvCM, varies dramatically throughout the course of evolution.

Inspired by these results, we develop a computational framework for predicting the EvCMs of a metabolic network by combining ideas from flux balance analysis, optimization, and evolutionary theory. This framework relates the structure of an EvCM to the expected mutational spectrum of genes, allowing us to make connections with long-term evolution experiments. We compare our theoretical predictions to experimentally observed mutation counts in the Lenski lines (1) and identify signatures of EvCMs, suggesting that the genotype-to-phenotype map stemming from metabolism likely places strong constraints on evolutionary dynamics.

## Results

To simulate the evolutionary dynamics of metabolic networks, we consider a clonal population of cells with an effective populations size, *N*_*e*_, grown in a constant environment. We focus on the strong-selection, weak-mutation regime where a single mutant can arise in the population and either fixate or go extinct (Figure 1A, (2)). The probability of fixation depends on the genotypes at time *t* of the mutant, 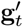, and parent, **g**_*t*_, through the selection coefficient *s*, defined as the difference in growth rates, *f*, between the mutant and parent strains: 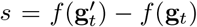. From population genetics, we know that the fixation probability is given by Kimura’s formula (14, 16):

**Fig. 1:**
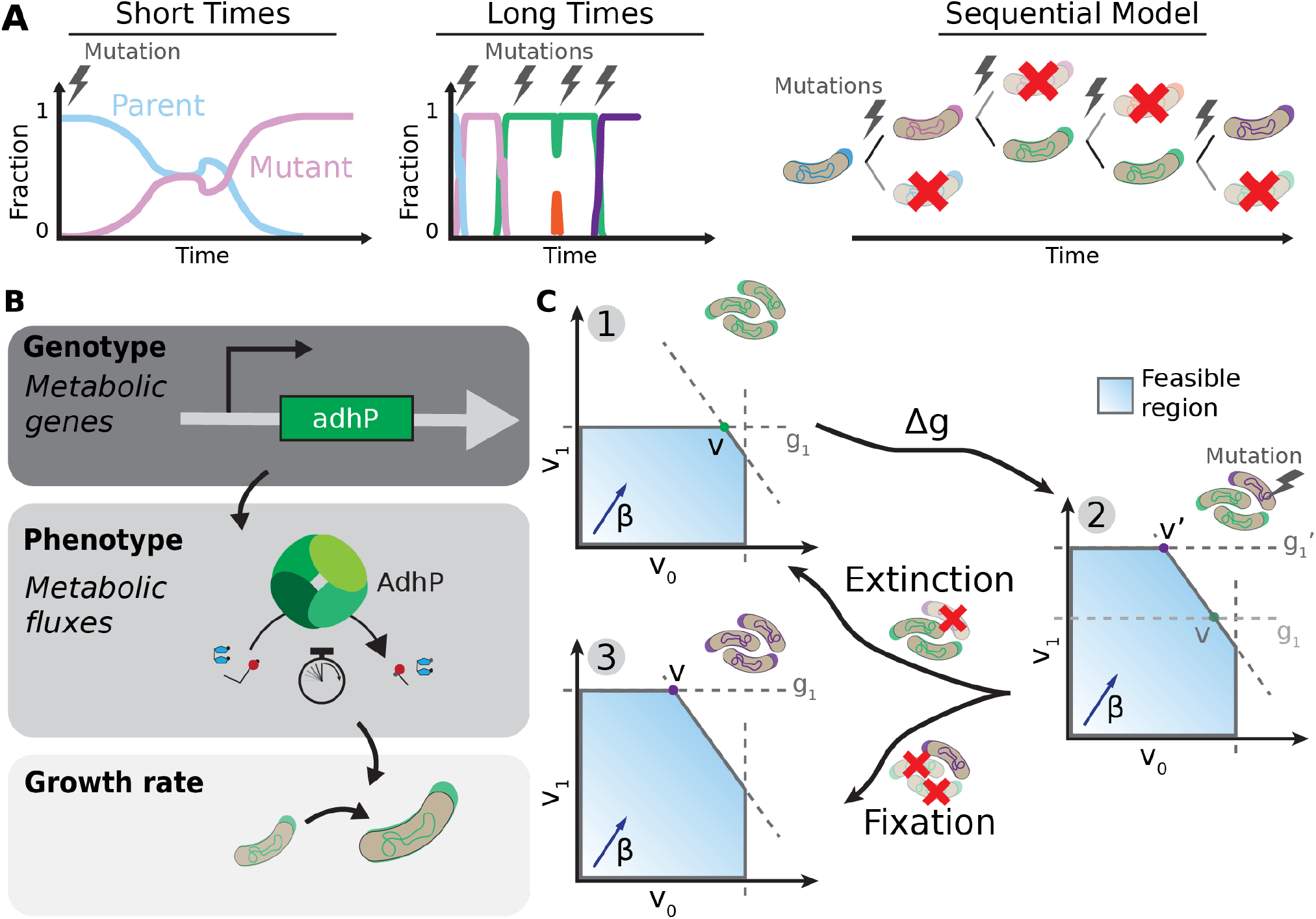
*(A)* Evolution can be modeled over various time scales from short to long. Over short times, a single mutation might occur, and the dynamics between the mutant and parent are important. At longer time scales, many mutations can occur. If the mutation rate is low, the population can reach a homogenous state between mutations; the mutant either overtakes the population or goes extinct. This logic allows us to model evolution as a sequential model where the frequency of populations is not tracked and instead the sequence of mutations is of interest. Kimura’s equation describes the probability of a new mutant fixing in this sequential limit. *(B)* Metabolism serves as a genotype-phenotype map. The genotype of interest is the metabolic genes. The phenotype of interest is the metabolic fluxes. Metabolic fluxes directly influence the growth rate, which can be used in Kimura’s equation. *(C)* A single step of evolutionary simulations. The initial set of genes **g** sets the phenotype **v**: The population is homogenous (1). A mutation affects one gene, **g**_1_, which changes its value to 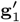. This mutation expands the feasible region of fluxes and a new set of fluxes **v**′ is achieved. Now, a single mutant is present in the population (2). Then, the mutant either fixes in the population (3) or goes extinct, leaving the parental population (1). As mentioned, the probability of fixation is determined by Kimura’s equation. Regardless of the outcome, the population is homogenous again.

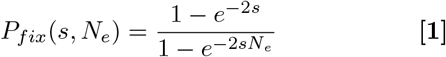

This expression arises from the interplay of stochastic (neutral) processes and selection. Selection dominates when the population size is large (|*s*| *>>* 1*/N*_*e*_) and neutral processes dominate with smaller populations (|*s*| *<<* 1*/N*_*e*_) (1, 17).

Calculating the selective advantage of a mutation requires a model of metabolism that explicitly relates genotype (genes) to phenotype (metabolic fluxes) and growth rate (Figure 1B). Here, we take advantage of the recent advances in metabolic modeling centered around flux balance analysis (FBA) (8, 9, 18). FBA is a constraint-based modeling approach that predicts metabolic fluxes — the rates at which metabolites flow through biochemical reactions in a metabolic network — by optimizing a biological objective function such as the growth rate. The method works by imposing stoichiometric constraints (mass balance: for internal metabolites, across all reactions the net production of the metabolite equals net consumption) and capacity constraints (bounds on reaction flux based on enzyme kinetics, thermodynamics, or the environment) on the flux distribution. Given these constraints, FBA uses linear programming to find the optimal flux distribution that maximizes the objective function, effectively predicting cellular metabolism without requiring detailed kinetic parameters for every reaction (9).

In mathematical terms, the most common formulation of FBA is a linear program of the form (9)

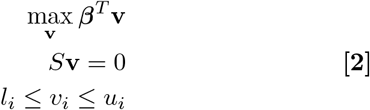

where **v** is a vector of metabolic fluxes with each element in the vector representing the flux through a single metabolic reaction in the network, *f* = *β*^*T*^ **v** is the flux-dependent growth rate, and *β* is a vector encoding the contribution of each reaction to growth. Mass conservation of metabolites is imposed through constraints on allowed flux configurations of the form *S***v** = 0, where *S* is the stoichiometric matrix of the metabolic network, which encodes how metabolites are produced and consumed in each reaction. Additional constraints, such as limits on influx of metabolites from the environment and the maximum allowable flux through a reaction due to genetic constraints, are encoded as inequalities, represented generically as *l*_*i*_ ≤ *v*_*i*_ ≤ *u*_*i*_.

One major shortcoming of this formulation as a model for metabolism is that there is no explicit dependence of the metabolic fluxes on the genotype. Instead, the dependence on genotype is implicit in the values of the lower- and upper-bound inequality constraints. For example, if an organism is missing the gene that encodes for an enzyme necessary for a particular metabolic reaction, then both the upper and lower bounds for the corresponding metabolic flux are set to zero (9). To rectify this, we draw inspiration from recent FBA models that study the effects of gene knockouts on metabolism by explicitly modeling the effect of genes on metabolism as inequality constraints on allowed metabolic flux configurations (9, 19– 21). We assume that the effect of a gene on its associated reaction is to limit the flux, rather than directly set the flux. Fluxes can also be limited due to non-genetic factors. For example, an exchange flux that governs how much of a metabolite is imported may be limited by metabolite concentration in the environment rather than gene expression. Furthermore, in the process of evolution, the constraint on a metabolic flux may shift from being limited by its genotype to the environment.

These considerations lead us to propose the following formulation of FBA, which we take as our model of metabolism:

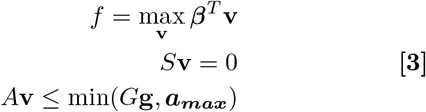

Notice that the inequality constraints now explicitly depend on the genotype of a cell, **g**, through matrices *A* and *G*, which allow for relationships between genes and reactions that are not one-to-one. For example, if two genes catalyze the same metabolic reaction, the reaction may be limited by the sum of the maximum allowable fluxes associated with the two genes (*i*.*e*., *v*_*i*_ ≤ *g*_*a*_ + *g*_*b*_). Similarly, multiple genes might need to cooperate to catalyze a reaction. In this case, the flux will be limited by the minimum of the genes’ maximum allowable fluxes. This relationship can be encoded in *A* (*i*.*e*., *v*_*i*_ ≤ *g*_*a*_ and *v*_*i*_ ≤ *g*_*b*_). We have also introduced a vector ***a***_***max***_ that encodes non-genetic limitations on metabolic fluxes. In most examples, we will assume that genetic limitations are the primary constraint on fluxes and take ***a***_***max***_ to be infinite. However, the immutable constraints arising from environmental limitations encoded by ***a***_***max***_ will play an important role in the evolutionary dynamics when examining realistic metabolic networks. Finally, the optimal growth rate *f* (**g**) is an explicit function of genotype through the constraints on metabolic fluxes.

To perform evolutionary simulations (Figure 1C, Methods) of a metabolic network described by *S, A, G*, and *β*, we start with a randomly selected value for the genes, **g**, and solve Equation 3 to obtain the phenotype, **v**. Then, we mutate **g** by adding random effect Δ**g** (typically a sparse vector altering a single gene) and resolving Equation 3 with **g** + Δ**g** to obtain the mutant phenotype **v**′. We calculate *s* = *β*^*T*^ **v**′ − *β*^*T*^ **v** and then allow the mutant to randomly fixate or go extinct with probability specified by Kimura’s equation (Equation 1). This process is repeated many times to simulate an evolutionary trajectory.

### A toy metabolic network for building intuition

Before considering large, realistic metabolic networks, it is useful to begin our discussion by considering the “toy metabolic network” shown in Figure 2A, consisting of four reactions and two metabolites. Despite its simplicity, this network contains many of the interesting behaviors seen in the evolution of more realistic metabolic models. For this reason, the toy network is extremely useful for building intuition about how evolutionary dynamics are shaped by genotype-to-phenotype maps.

**Fig. 2:**
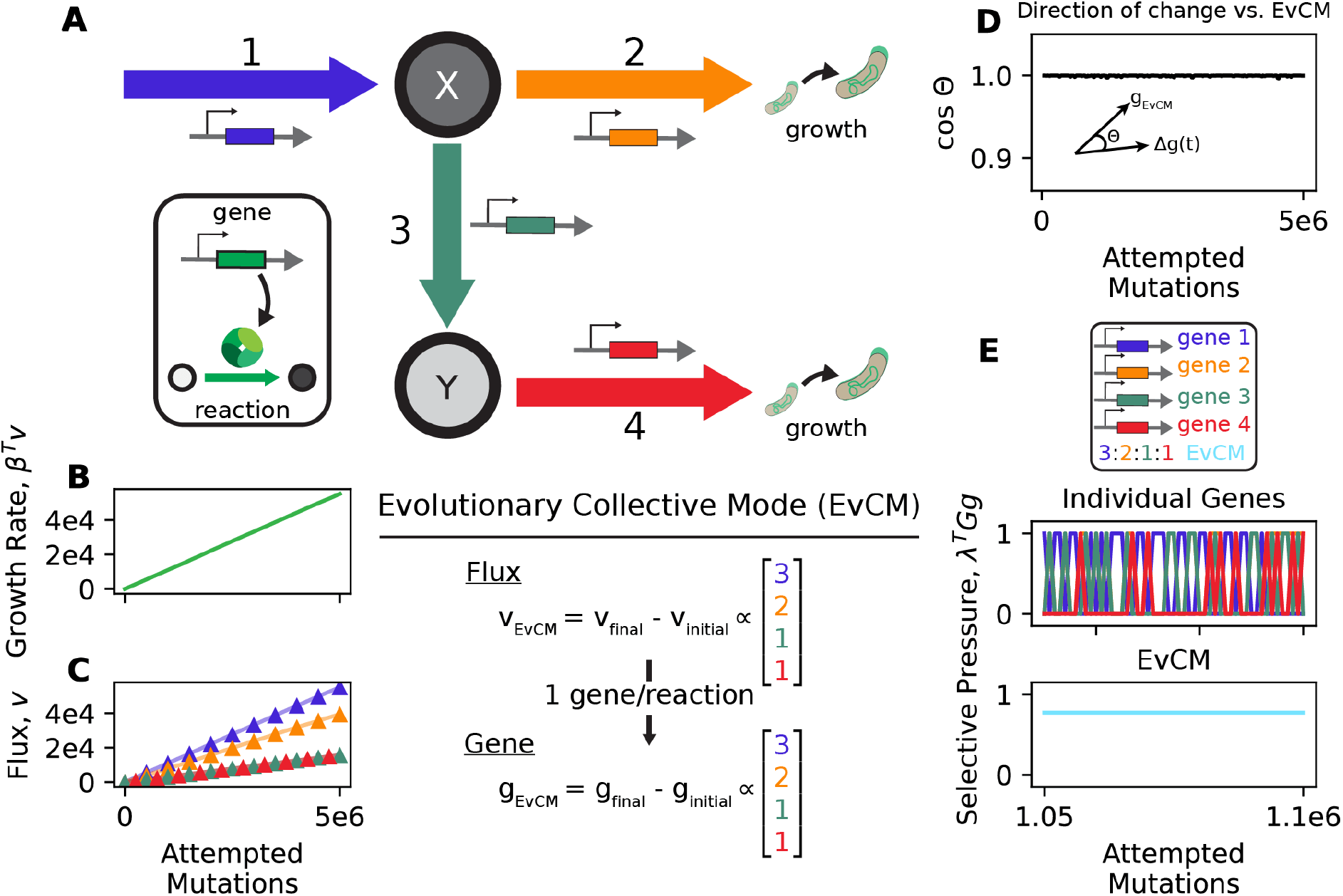
*(A)* Diagram of the toy metabolic network. This network has two metabolites, *X* and *Y*. *X* can be imported into the cell (reaction 1) and consumed for growth (reaction 2). *X* can also be converted into *Y* (reaction 3), which can also be consumed for growth (reaction 4). Each reaction has one associated gene that undergoes evolution. During evolutionary simulation, the growth rate *(B)* and fluxes *(C)* of the toy metabolic network increase as more mutations are attempted. Using the genes, we can calculate the evolutionary collective mode (EvCM) as the change in genes throughout evolution. We confirmed that evolution is proceeding along the EvCM by examining the cosine similarity between the estimated EvCM (**g**_*EvCM*_) and the local change in genes. The cosine similarity is high, indicating that the vectors are pointing in the same direction. *(E)* The selective pressure on individual genes and the EvCM were estimated over a small window of the evolutionary simulation. The selective pressure on the EvCM is nearly constant, while the selective pressure on the individual genes varies consistently across time.

In the toy network, a metabolite *X* is imported into the cell (reaction 1) and can either be consumed (reaction 2) or converted into a metabolite *Y* (reaction 3), which is also consumed (reaction 4). The growth rate of the cell is the sum of fluxes through reactions 2 and 4. We also assume a simple genotype-to-phenotype map where each of the four reactions is regulated by a different gene that sets an upper bound for the flux through the reaction (both *A* and *G* are identity matrix in Equation 3; SI Appendix Section 4A).

### Evolutionary simulations show the emergence of an Evolutionary Collective Mode

We begin by considering the case where the metabolite *X* is in excess in the environment and hence all metabolic constraints on fluxes are genetic (*i*.*e*., ***a***_***max***_ is infinite; we extend below to the case where *X* is limited in the environment). We simulated the evolution of this toy network and the resulting evolutionary dynamics are shown in Figure 2. As mutations accumulate, the growth rate of the population increases with time (Figure 2B), the population gets fitter, and the flux through all four reactions increases (Figure 2C), as does the upper-bound constraint on the fluxes encoded by the four genes (SI Appendix Figure S1). Since the external metabolite *X* is always in excess, evolution proceeds indefinitely with continual fitness gains since mutations can always increase the total flux of nutrients into the cell.

One striking aspect of these results is that *the ratio of the fluxes through each of the four reactions, as well as the ratio of the upper-bound constraints* on each of the four reactions, is approximately constant and fixed to a ratio of 3 : 2 : 1 : 1 (Figure 2D, SI Appendix Figure S1). This indicates that the evolutionary dynamics have a preferred evolutionary direction in both genetic (constraint) and phenotypic space. A natural question one can ask is: how reproducible is this observation across replicates?

Even for this simple toy network, the answer is far from obvious. There is a huge degeneracy in flux space resulting from the fact that the growth rate is just the sum of the fluxes through reactions 2 and 4. For example, routing all the flux from reaction 1 through reactions 3 and 4 results in the same growth rate as routing all the flux from reaction 1 through reaction 2. As a result, it is difficult to predict *a priori* how the system will evolve if we “replay the tape of evolution” (22) with different sequences of random mutation.

To answer this question, we performed five replicate simulations of evolution in the toy network. We found that, despite the degeneracy discussed above, in all five replicates, both the fluxes and the upper-bound constraints associated with each of the four reactions evolve in an approximately constant 3:2:1:1 ratio (SI Appendix Figure S2). Since in this toy network both *A* and *G* are the identity matrix and ***a***_***max***_ is infinite, we can directly identify the upper-bound constraints with the genotype vector **g**. Our simulations indicate that evolution proceeds along a preferred direction in this constraint/gene space, namely in the direction of the vector **g**_*EvCM*_ = [3, 2, 1, 1]. We call this preferred direction the Evolutionary Collective Mode (EvCM). The existence of the EvCM reflects constraints placed on the evolutionary dynamics by the genotype-to-phenotype map resulting from metabolism. Even though each of the four genes in our network independently regulate a different reaction, they become coupled in the evolutionary dynamics so that mutations occur at a fixed ratio.

### The EvCM corresponds to directions of high, constant selective pressure

To gain more insight into why the EvCM emerges in our evolutionary simulations, it is helpful to think about the selective pressure on genes. Consider a population with genotype **g**^∗^ that undergoes a mutation (Δ**g**), resulting in a change in the growth rate

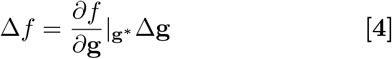

We call 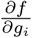 the selective pressure on gene *i*. Note that in general, the selective pressure will depend strongly on the current genotype **g**^∗^ since the effect of a mutation depends on the genetic background through the upperbound constraints on fluxes. For this reason, we expect that the selective pressure on individual genes will vary significantly across the evolutionary dynamics. As shown in Figure 2E, this is indeed the case for all four genes in our toy metabolic network.

Despite this variability, our simulations also show that the growth rate of the population increases approximately linearly with evolutionary time (Figure 2B). These two observations can only be reconciled if the time-averaged selective advantage across all fixed mutations (the change in growth rate averaged over mutations across all genes in a time window) is approximately constant throughout the evolutionary dynamics. Since we have already observed that evolution proceeds preferentially along the EvCM, **g**_*EvCM*_, this suggests that the selective pressure along the EvCM, defined as 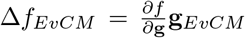, must be consistently high and constant. Surprisingly, this must be true even though the selective pressure on the individual genes that make up the EvCM fluctuates dramatically across the evolutionary dynamics.

To test if this is indeed the case, we used our evolutionary simulations to calculate the selective pressure on individual genes and in the direction of the EvCM as a function of time. To do so, we exploited the fact that the selective pressure is closely related to the shadow prices associated with metabolic constraints and thus can be calculated using the dual formulation of the linear programming objective in Equation 3 (Methods, (23)). As shown in Figure 2D&E, the selective pressure on the direction of the EvCM is consistently high and constant, independent of the population’s current genotype and evolutionary history. These simulations suggest that the EvCM corresponds to directions in genotype space with high, constant selective pressure. We show below that these ideas can also be used to computationally predict the collective modes of any metabolic network.

### Non-genetic constraints lead to evolutionary regimes

Thus far, in all simulations we have assumed that mutations can always increase the flux through any reaction (*i*.*e*., ***a***_***max***_ is infinite in Equation 3). However, we expect that in most evolutionary scenarios some reactions will face immutable constraints. This limit can arise due to environmental limitations (*e*.*g*., the external metabolite X in Figure 2A is at a low concentration) or due to an inherent limit on expression levels of proteins.

To understand how the presence of these non-genetic constraints affects evolutionary dynamics, we created two additional toy networks (Figure 3). In the first network, there is a maximum allowable exchange flux through reaction 1, mimicking evolution in an environment where extracellular resources are limited. We assume that the *a*_*max*_ for reaction 1 is sufficiently small so that the upperbound constraint on reaction 1 is fixed and its gene can no longer evolve. In the second network, there is a maximum allowable flux through reaction 3, mimicking a scenario where the enzyme for reaction 3 cannot be produced at higher concentrations. We again assume that the *a*_*max*_ for this reaction is sufficiently small that gene 3 can no longer evolve.

**Fig. 3:**
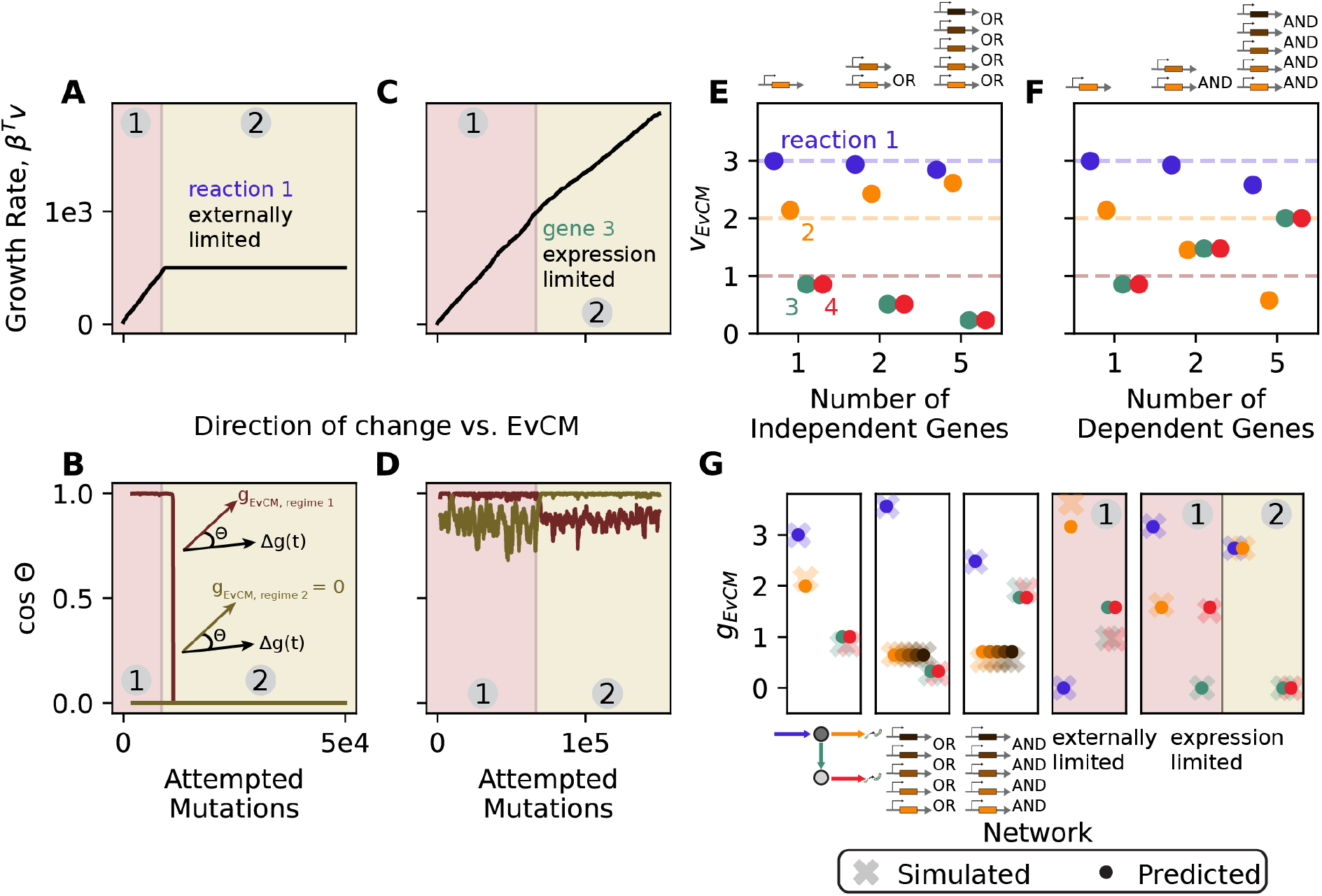
Variations of the toy network reveal how resource limitations and gene-reaction relationships shape the EvCM. *(A)* Simulations were performed where reaction 1 is externally limited (*a*_*max*_ ≤ ∞ for reaction 1). Initially, the growth rate increases. However, once the externally limited flux is reached, evolution stops, splitting the simulation into two evolutionary regimes. *(B)* The cosine similarity between direction of change and the EvCM for each regime. The EvCM for the second regime is 0, so the cosine similarity is 0. As the first regime ends and the second starts, the direction of evolution becomes less aligned with the EvCM of the first regime. *(C)* Simulations were performed where reaction 3 is limited by the expression of its associated gene (*a*_*max*_ ≤ ∞ for reaction 3). Again, evolution is split into two regimes. However, the reaction 2 pathway can still evolve, so growth rate continues to increase. *(D)* The cosine similarity between direction of change and the EvCM for each regime. As the first regime ends and the second starts, the direction of evolution becomes less aligned with the EvCM of the first regime and more aligned with the EvCM of the second regime. *(E)* **v**_*EvCM*_ as multiple genes can independently increase the maximum allowable flux of reaction 2 (OR relationship). As more genes are added, the EvCM favors flux through reaction 2 over the reaction 3/4 pathway. *(F)* **v**_*EvCM*_ as multiple genes dependently increase the maximum allowable flux of reaction 2 (AND relationship). As more genes are added, the EvCM favors flux through the reaction 3/4 pathway over reaction 2. *(G)* Predictions of the EvCM compared to simulation EvCM for metabolic networks discussed to this point. Predictions occur at the level of genes and in terms of the maximum allowable flux.

The evolutionary dynamics of these networks are shown in Figure 3. Initially, both these networks evolve along an EvCM, as in the original toy metabolic network. For both networks, the growth rate increases approximately linearly in time with a fixed slope. However, the presence of immutable non-genetic constraints dramatically changes the evolutionary dynamics of both networks once the maximum allowable flux is reached through the clamped reaction. In the first network, evolution stops entirely (the growth rate plateaus). This is because cells can no longer increase the amount of nutrients they import, making it impossible to keep increasing their growth rate.

The evolutionary dynamics of the second network are much more interesting. In this case, the evolution continues but along a new EvCM where the growth rate also grows linearly in time, but now with a different slope. In this regime, the ratio of fluxes and upperbound constraints are again constant, but with different numerical values from the first period. We call this behavior where the evolution shifts between different EvCMs the emergence of distinct evolutionary regimes.

To test whether more than two evolutionary regimes can arise during evolution, we performed additional simulations (SI Appendix Section 4B). SI Appendix Figure S3 shows the evolutionary dynamics for the first network (reaction 1 cannot evolve) if we modify the fitness function so that the flux in reaction 2 contributes to growth slightly less than the flux through reaction 4. In this case, we see that there are three distinct evolutionary regimes: an initial EvCM, which transitions to a second EvCM, before evolution stops. Additional evolutionary regimes can emerge if we make our toy network more complicated. For example, if we take the same network and now add a third pathway that contributes to growth (SI Appendix Figure S3), we find that are now four evolutionary regimes. The system now transition between three distinct EvCM before evolution stops.

The emergence of distinct evolutionary regimes reflects that fact that as reactions can no longer evolve, the optimal direction for evolution changes, resulting in the evolutionary dynamics following a new EvCMs. For this reason, we expect evolutionary regimes to emerge in any realistic or sufficiently complex metabolic network.

### Evolvability of reactions shapes the structure of the EvCM

Thus far we have said very little about why the EvCM of our toy network points in the direction it does. What gives rise to the conserved 3:2:1:1 ratio for fluxes and upper-bound constraints seen in our evolutionary simulations? Here, we present evidence suggesting that the EvCM reflects an interplay between biophysical constraints stemming from mass conservation and the “evolvability” of reactions, with reactions that are easier to evolve contributing more to the collective mode. Consider the flux through each reaction in the EvCM. Examining Figure 2A, it is clear that mass balance requires that the flux through reaction 1 must equal the total flux through reactions 2 and 3. Indeed, looking at the EvCM **v**_*EvCM*_ = **g**_*EvCM*_ = [3, 2, 1, 1], we see that the total flux through reaction 1 (*v*_1_ = 3) equals the sum of the flux through reaction 2 (*v*_2_ = 2) and reaction 3 (*v*_3_ = 1). Similarly, the total flux through reaction 3 must equal the flux through reaction 4. However, the 2:1 ratio between the metabolic fluxes through reactions 2 and 3 cannot be explained by mass balance constraints. This suggests the structure of the EvCM also depends on mapping between genes and fluxes/constraints (the genotype-to-phenotype map).

Recall that the growth rate of a cell in our toy network is the sum of the fluxes through reaction 2 and reaction 4. Looking at the structure of our toy network in Figure 2, it is clear that the cell can increase its growth rate in two distinct ways: either by increasing the flux through reaction 2 or increasing the flux through reactions 3 and 4. A single beneficial mutation to gene 2, which sets the upper bound on the flux through reaction 2, is sufficient to increase the flux through reaction 2. In contrast, the flux through reaction 4 is limited by the minimum of upperbound constraints of reactions 3 and reaction 4. For this reason, on average, it takes twice as many mutations to increase fitness using reaction 3 and 4 compared to reaction 2; both reactions 3 and 4 will need to be mutated to increase the flux through this pathway. This suggests that mutations should fix twice as often in the gene for reaction 2 compared to genes for reactions 3 or 4. Since all genes are equally likely to mutate, this suggests that the maximum allowable flux (and upper-bound constraint) of reaction 2 should grow twice as fast as those of reactions 3 and 4 during the course of evolution.

This analysis suggests that the structure of the metabolic network influences evolvability and the collective mode. If more genes must be mutated for the same fitness benefit, the associated portion of the metabolic network will have smaller contributions to the collective mode. To test this hypothesis, we performed further simulations where we added additional genes to our toy metabolic network. First, we considered a scenario where multiple genes can independently catalyze the same reaction (*i*.*e*., independently increase the upper-bound constraint on the corresponding reaction). For example, in *E. coli*, the isomerization of 3-phosphoglycerate into 2-phosphoglycerate can be catalyzed by gpmM or gpmA (24). When multiple genes are present, we expect that a reaction should be more evolvable, and thus, a mutation that increases the maximum allowable flux for the reaction should be more likely to occur.

To computationally test this, we augmented the toy network by adding additional genes that code for reaction 2 (Figure 3E). We assume genes can independently catalyze the reaction, so the maximum allowable flux for the reaction is the sum of the maximum allowable fluxes associated with each gene. When we examine the collective mode of these metabolic networks, we see that the relative flux through reaction 2 increases as more genes can catalyze it, while the flux through reactions 3 and 4 necessarily decreases (Figure 3E).

A second gene-reaction relationship of interest is when multiple distinct genes must coordinate to catalyze a reaction. For example, in *E. coli*, the conversion of succinyl-CoA to coenzyme A and succinate in the citric acid cycle requires sucD and sucC, the alpha and beta subunits of succinyl-CoA synthetase, respectively (25). In this setting, we expected the evolvability of the reaction to decrease. If we assume that the subunits must cooperate stoichiometrically, increasing the maximum allowable flux with one gene will not necessarily lead to an increase in the maximum allowable flux of the reaction.

To test this hypothesis, we again augmented the toy network. We assumed that the protein catalyzing reaction 2 had multiple subunits with distinct genes (Figure 3F). Because the genes must cooperate to catalyze the reaction, we assume the maximum allowable flux of the reaction is the minimum of the maximum allowable fluxes associated with the genes. When we examine the collective mode of these metabolic networks, we see that the relative flux through reaction 2 decreases as more subunits are needed to catalyze it, while the flux through reactions 3 and 4 increases (Figure 3F).

Taken together, these results indicate that the structure of the EvCM results from a delicate interplay between biophysical constraints and how evolvable a reaction is. When multiple genes catalyze a reaction independently, evolvability increases, and the reaction is favored in the collective mode. Alternatively, when multiple genes are required, evolvability decreases, and the reaction is disfavored in the collective mode.

### Non-identifiability of genes

With a one-to-one mapping between genes and reactions the EvCM could be easily mapped between genotype and phenotype, but with multiple genes associated with a given reaction it is less clear if a single, preferred direction in phenotype space would also be reflected in a single, preferred direction in genotype space. Mathematically, this might be highlighted when there is degeneracy in the *G* matrix, as happens when multiple genes independently catalyze a reaction. To examine this, we looked at the variability of the change in genotype compared with the variability of the change in the associated reaction fluxes over relatively short periods of evolution (5 *×* 10^4^ attempted mutations). We found that when a gene uniquely maps to a reaction, the variability is similar. However, when multiple genes independently contribute to a single reaction, there is greater variability at the level of genes than at the level of fluxes (SI Appendix Figure S4). This result suggests that evolution cannot differentiate among multiple independently contributing genes in this setting, and that the evolutionary dynamics converge to the EvCM as a single direction in phenotypic, but not necessarily genotypic, space.

### Predicting the EvCM from the genotype to phenotype map

We saw above that the EvCM has two key properties – high and constant selective pressure – and that we could make sense of the “preference” for this direction through the notion of evolvability of reactions. Now, we ask if we can formalize these concepts with the goal of predicting the direction of evolution for a given metabolic network. We approach this problem first in the case of a single evolutionary regime, then discuss prediction when non-genetic constraints lead to multiple regimes.

The shadow prices introduced above play a central role in prediction. Recall that shadow prices capture the selective pressure on each constraint and that they shift as the network evolves and thus are a function of genotype; at time *t* we have shadow prices **λ**(**g**_*t*_). These are mapped to selective pressure on genotype through the *G* matrix. The selective pressure on a direction of genotypic change **g**_*EvCM*_ can then be written as *p*(**g**_*EvCM*_, **g**_*t*_) = **λ**(**g**_*t*_)^*T*^ *G***g**_*EvCM*_. With the property of constant selective pressure in mind, we’re interested in vectors **g**_*EvCM*_ for which *p*(**g**_*EvCM*_, **g**_*t*_) does not vary, even as the genotype **g**_*t*_ changes.

At first glance, it seems challenging to identify such a vector since the genotype continuously evolves (especially in the absence of non-genetic constraints), and it is therefore difficult to say over what set of genotypes the selective pressure should remain constant. However, even when the genotype continues to evolve, the genotype-to-phenotype map defines a limited set of shadow price vectors; that is, the space of selective pressure is limited even when the genotype is not.

For instance, in the toy network of Figure 2A, there are only three possible values of λ (SI Appendix Figure S5). These reflect the different possibilities for limiting reactions under a mass balance constraint: when the import (reaction 1) is limiting, there is selective pressure on that reaction and no other. If the import reaction is not limiting, then there are two scenarios reflecting pressure to increase downstream flux, reflecting whether reaction 3 or 4 is limiting in the bottom branch (SI Appendix Figure S5). Thus, in this case a direction **g**_*EvCM*_ with constant pressure over just these three values of **λ** will have constant pressure over all genotypes and timepoints.

In principle, there could be many vectors with the property of constant selective pressure (in the toy network this includes all vectors that, when interpreted as reaction fluxes, achieve mass balance, though the result is more complicated with non-identity *A* or *G*), and so to predict the EvCM we need a way to differentiate among this set. To do so, we posit that among this set the EvCM will have the maximum selective pressure. We formalize this as a constrained optimization, searching for the direction of maximal selective pressure, subject to the constraint that this pressure is constant across all values of **λ** (SI Appendix Section 6). Given the fitness function, *S, G, A*, and *β*, we can enumerate the **λ**s, then solve a quadratic program to arrive at a predicted direction of evolution.

For the toy network, this routine accurately predicts the EvCMs. For the simple case in Figure 2, the prediction is **g**_*EvCM*_ = [3, 2, 1, 1], matching simulation outcomes (Figure 3G). Similarly, we can accurately predict how the EvCM changes as independent or dependent genes are added for reaction 2. When multiple independent genes are added, **v**_*EvCM*_ favors reaction 2, while when dependent genes are added, **v**_*EvCM*_ disfavors reaction 2, consistent with our simulation results.

With non-genetic constraints and different evolutionary regimes, we can also accurately predict the EvCM. To incorporate the effect of non-genetic constraints, we force the non-evolvable genes to be 0 in **g**_*EvCM*_. Then, we still have that **g**_*EvCM*_ has constant selective pressure over the values of λ reached by evolution (SI Appendix Section 6D). When we use this approach in the presence of regimes, we can accurately predict the EvCM within each regime (Figure 3G).

### EvCMs in random metabolic networks

We next examined whether properties of the toy network – including the presence of an EvCM and our ability to predict the EvCM – generalize to larger, randomly generated metabolic networks. We examined 400 networks with 25-100 metabolites, 50-200 reactions, a randomly generated stoichiometric matrix, and random biomass reactions, while varying gene-reaction relationships from one-to-one to more complex (with non-identity *A* or *G*; SI Appendix Section 7).

Two small differences from simulations with the toy network are worth pointing out. Positive and negative (reverse direction) fluxes may appear (the formalism introduced above handles this transparently), and conceivably a randomly generated network could fail to evolve. We avoid this scenario in our random selection of metabolic networks (SI Appendix Section 7A) and we focus our analysis here on evolving networks, in which mutations accumulate, the population gets fitter, and where the magnitude of constraints and flux increases.

In the evolving random networks, we consistently observe EvCMs (Figure 4). First, the direction of evolution is constant. We find that, across a given simulation, the changes in flux or genes vary along a single direction (SI Appendix Figure S7). Moreover, the selective advantage of mutations in this direction appears to be constant. We observe this by noting that the population fitness over time can be described by a linear function (with *R*^2^ values *>*0.99 across all random simulations), and by observing that that the shadow price-based calculation of selective pressure on a given network’s direction of evolution is nearly constant (Figure 4B; SI Appendix Figure S6).

**Fig. 4:**
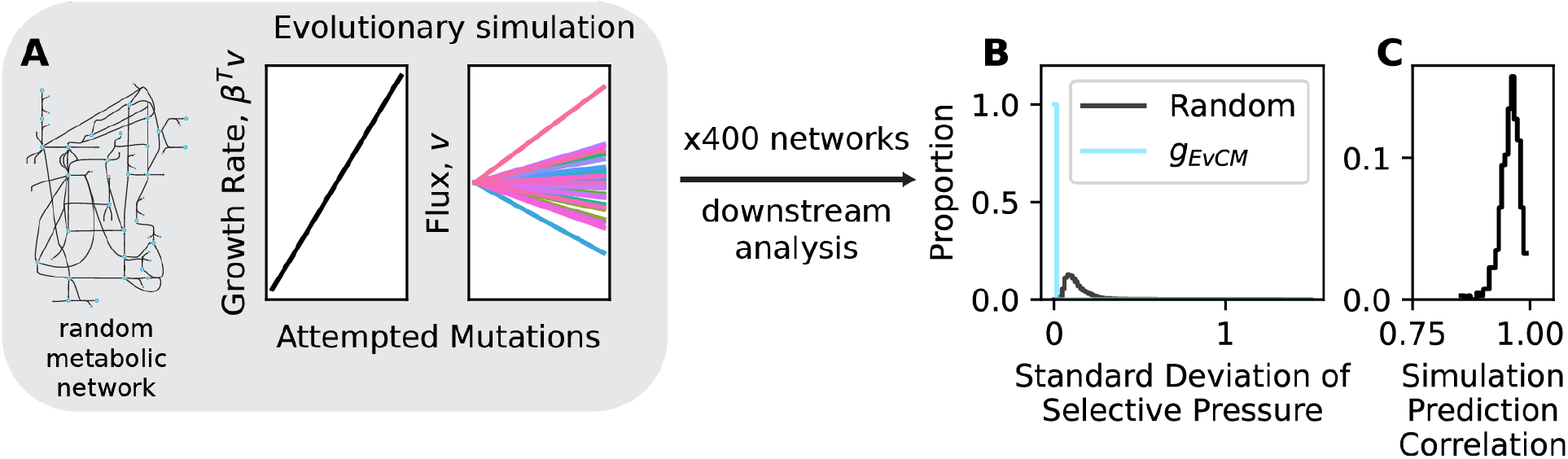
*(A)* Random metabolic networks were generated (as described in SI Appendix Section 7) and evolutionary simulations were performed. The growth rate and flux increased in magnitude for each simulation, indicating evolution is occurring. 400 random metabolic networks were simulated. *(B)* The distribution of the standard deviation of the selective pressure on each network’s EvCM compared to the standard deviation of the selective pressure on random directions for each network. The selective pressure on the EvCM is constant for each network. *(C)* The distribution of correlations between simulation EvCM and predicted EvCM using the criteria of constant and maximal selective pressure. The simulation EvCM can be accurately predicted.

Second, evolutionary outcomes are consistent across trials. Replicate trials of the same network produce the same outcome (SI Appendix Figure S7; *>*99% correlation of genes and fluxes), even though many combinations of fluxes could achieve the same final fitness. Thus, a given random network is always driven by the same EvCM.

Third, evolvability shapes the collective mode in random metabolic networks. As before, we find that when more genes can independently catalyze a reaction, there is greater evolvability, and an increasing representation of that reaction in the collective mode (SI Appendix Figure S8). When more dependent genes are required, there is less evolvability, and a decreasing representation in the collective mode (SI Appendix Figure S8). We also observe the effects of gene non-identifiability. We see that the difference in variability of fluxes and genes is positively correlated with the number of independent genes for a reaction, consistent with our earlier results in the toy network (SI Appendix Figure S8). This result again suggests that evolution cannot differentiate among multiple genes that independently contribute to the same constraint.

Finally, we used the procedure above to predict the collective mode of each randomly generated metabolic network. With few exceptions, these predictions are *>*90% correlated with simulation outcomes at the gene level (SI Appendix Figure S13).

### EvCMs of E. coli central metabolism

We next simulated the evolution of a more realistic network: *E. coli* central metabolism (Figure 5A). We use the ‘e coli core’ model from BiGG Models (18) with slight adjustments (SI Appendix Section 8). This network has 72 metabolites and 95 reactions. The reactions primarily consist of glycolysis, the citric acid cycle, and oxidative phosphorylation. Additionally, each reaction has an associated ‘gene rule’ that relates genes to reactions, leading to 137 genes in the model. We used these gene rules to generate *A* and *G* matrices (SI Appendix Section 2). This model assumes that growth is limited by glucose and no other energy sources can be imported into the cell (18). As a result, there is a non-genetic constraint defining a fixed maximum allowable flux on the exchange reaction for glucose.

**Fig. 5:**
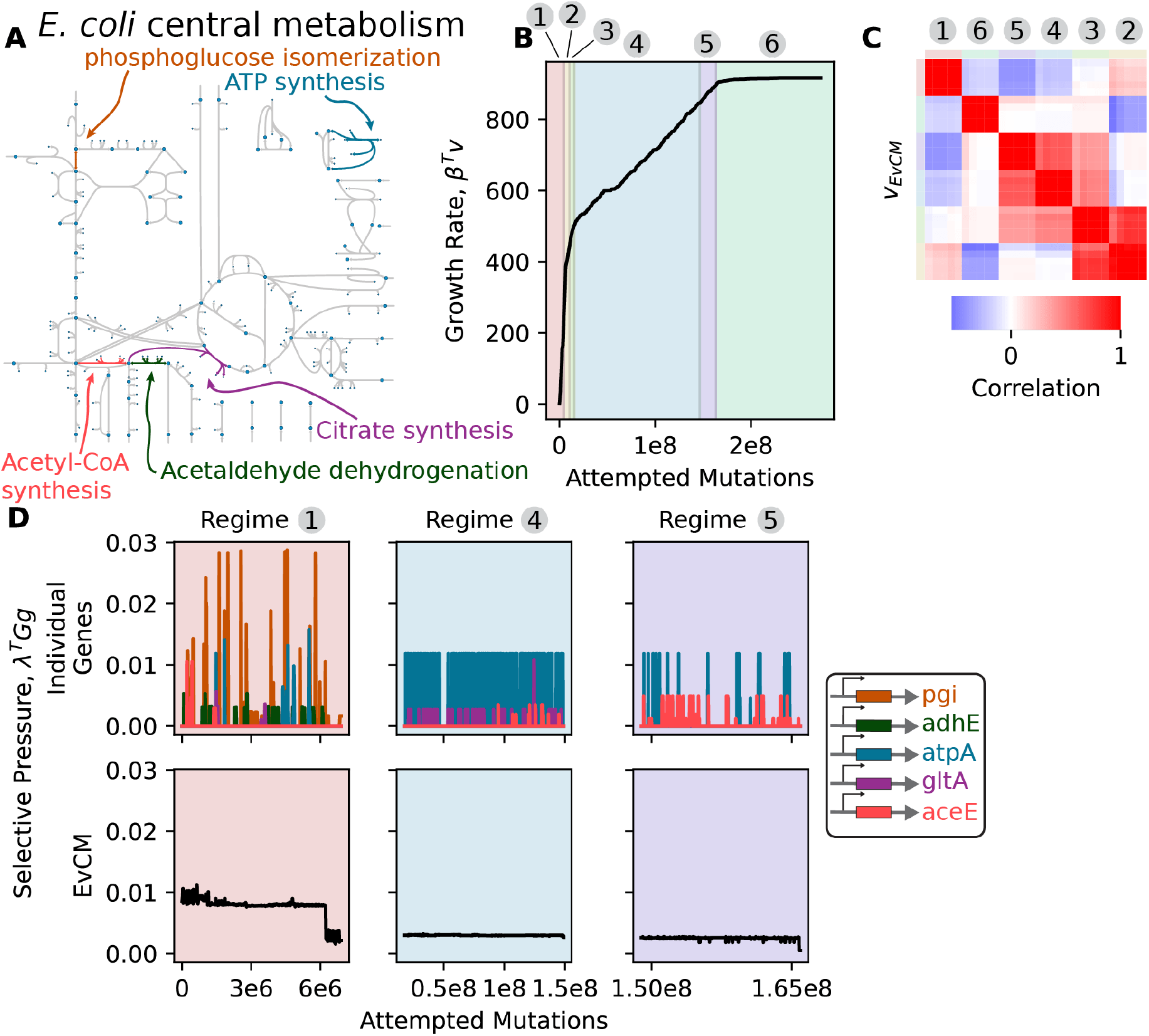
Simulations of the *E. coli* central metabolism have evolutionary regimes with EvCMs. *(A*) The *E. coli* central metabolism broadly consists of the glycolysis pathway, citric acid cycle, and oxidative phosphorylation. The network consists of 95 reactions, but five reactions are highlighted to capture the broad changes in the network. *(B)* The growth rate of the metabolic network increases as more mutations are attempted. The simulation has six evolutionary regimes, which can be observed partially from changes in the slope of the growth rate (see SI Appendix Figure S10 for replicates). *(C)* Five replicate simulations were performed, each of which had six evolutionary regimes. The EvCM at the flux level was estimated for each regime, and the correlations of all EvCMs were calculated. Hierarchical clustering was performed on each EvCM, and the EvCMs cluster by regime, as indicated by color along border, showing consistency across replicates. *(D)* The selective pressure was calculated on individual genes (associated with reactions highlighted in *(A)*) and the EvCM within regimes 1, 4, and 5. The selective pressure on the individual genes is variable across time, and often zero. In contrast, the selective pressure on the EvCM is nearly constant across time and always greater than zero.

We analyzed the results of 5 simulation trials with up to ∼ 2 *×* 10^8^ attempted mutation. Multiple regimes are evident as periods with different slopes in fitness over time for a given trial (Figure 5B). We found six regimes based on abrupt changes in flux directions (SI Appendix Figure S10), with distinct directions of evolution within each regime. The direction of evolution within each regime and the ordering of regimes was strikingly consistent across replicate trials (SI Appendix Figures S10). Within each regime the direction of evolution evaluated as the ratio of fluxes was constant (SI Appendix Figure S10), as was the shadow price-based selective pressure on these directions (Figure 5, SI Appendix Figure S10). Predictions of the collective mode in each regime at the gene level were *>*80% correlated with simulation outcomes (SI Appendix Figure S9). In aggregate, these results show that in simulation the central metabolism of *E. coli* evolves along collective modes.

While this simulation is only an approximation of evolution, we share what occurs biologically in each regime. Identifying specific regimes is challenging, so we speak in broad terms and only focus on the largest and most obvious features. Initially, the flux through glycolysis increases and ethanol is fermented. After a transition period, the largest regime (labeled 4), appears to focus on utilizing the citric acid cycle and oxidative phosphorylation to generate ATP. The last portion of the simulation appears to be further refinements, including switching from using pyruvate formate lyase to pyruvate dehydrogenase for generation of acetyl-CoA. Again, we emphasize these collective modes are likely sensitive to what reactions and genes are included in the metabolic network. However, these results do match our expectations. The proteins in the steps of oxidative phosphorylation require coordination of many genes (26), while the steps of ethanol fermentation generally only have a single or few genes (27). As a result, from the perspective of our previous results, the fermentation pathway is more evolvable than oxidative phosphorylation and should be favored by the EvCM initially. Oxidative phosphorylation is evolved once growth rate can no longer be increased with fermentation.

### Evidence of EvCMs in a long-term evolution experiment

Finally, we turn to experimental data and look for evidence of collective modes in the Lenski lines. The Lenski lines are a long-term evolution experiment where *E. coli* has been serially cultured for 60,000 generations (1). Twelve initially identical lines have been independently cultured, and fitness increased in all lines throughout the experiment. Six of the lines developed a hypermutator phenotype (1, 28), which makes analysis difficult, so we focus on the normal mutators. While metabolic fluxes (and constraints) were not systematically measured, the structure of EvCMs should be reflected in the mutational spectrum across genes. Metagenomic sequencing has been performed every 500 generations (1), and we looked for evidence of EvCMs in this dataset.

The normal mutator lines display a remarkable consistency in evolutionary dynamics and outcomes. Each of the lines achieved rapid fitness gains during the first ∼10,000 generations, followed by a period of slower, but still accumulating fitness gains for the duration of the experiment (Figure 6A). This is reminiscent of a two regime outcome seen in simulation (Figure 3C). Further, the mutational spectra (counts of fixed mutations per gene across all 3,969 genes mutated in the experiment) accumulated by the end of the experiment are 25% correlated on average (SI Appendix Figure S11).

**Fig. 6:**
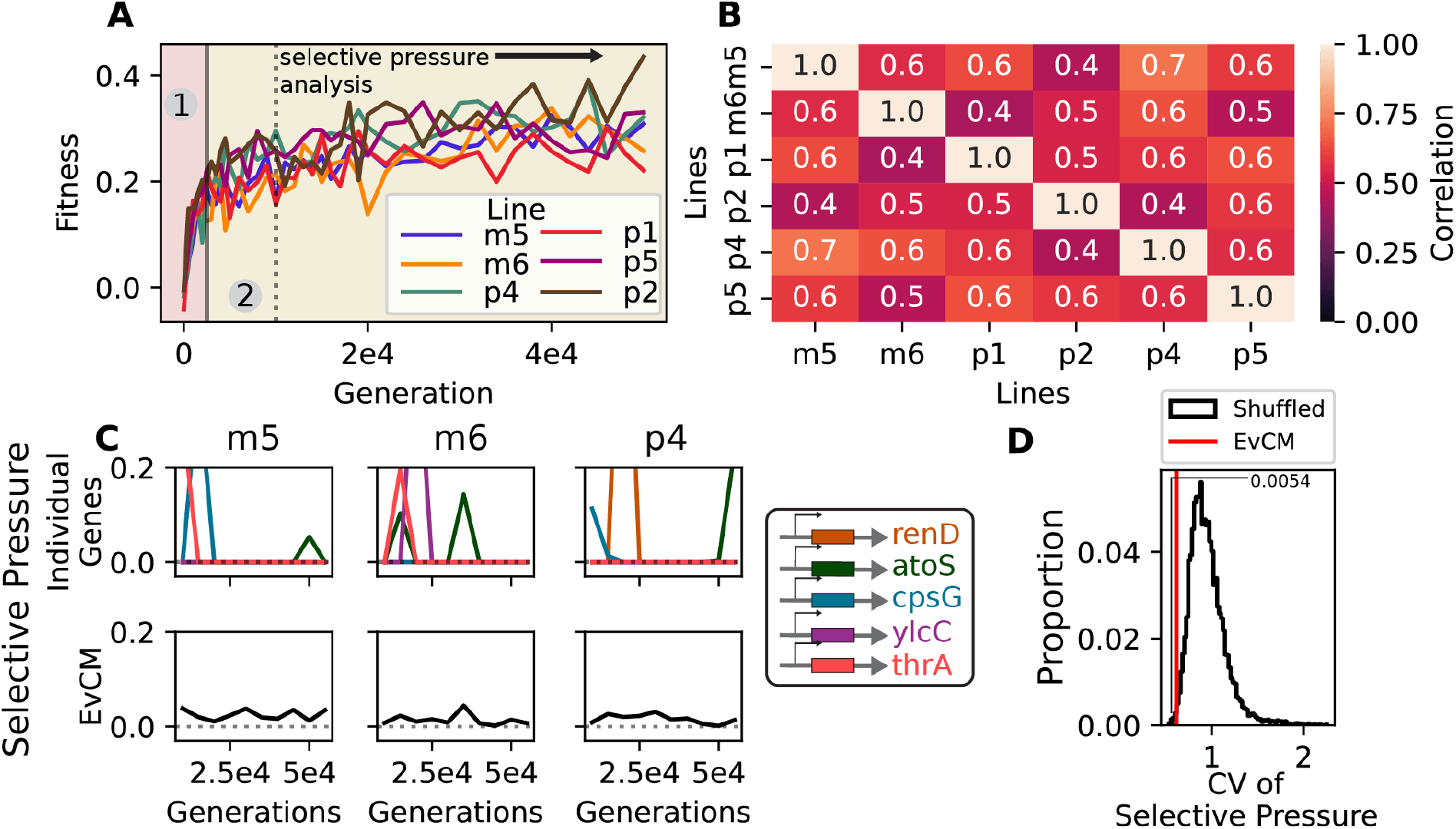
The Lenski lines were analyzed to look for evidence of EvCMs. *(A)* The fitness of the non-mutator lines over the course of the experiment. Within our framework, the fitness trajectories appear consistent with multiple regimes. The end of the first regime was estimated at 2,500 generations. *(B)* While the mutation counts across the entire experiment are weakly (SI Appendix Figure S11), regulatory-adjusted counted are more correlated across lines, suggesting the Lenski lines may be evolving in a similar manner. *(C)* The Lenski lines were analyzed for evidence of constant selective pressure. To avoid any regions of overlap between the regimes, only data from beyond generation 10,000 was analyzed (indicated in A). The selective pressure within the three held-out lines was calculated on individual genes and the estimated direction of evolution. The shown genes had the largest average selective pressure across times and lines. The selective pressure on the individual genes is variable across time, and often zero. In contrast, the selective pressure on the EvCM is nearly constant across time and greater than zero. *(D)* The coefficient of variation of the selective pressure on the estimated EvCM across lines and time compared to 10,000 samples from a null distribution. The null distribution was obtained by shuffling the estimated direction of evolution and recalculating the coefficient of variation of the selective pressure on it (SI Appendix Section 9B).

Since our previous analysis of non-identifiability of genes indicated that a common direction of fluxes can emerge even as directions of evolution vary at the level of genes, we wondered if there might be more in common among the lines than suggested by the baseline correlation. Notably, many of the most mutated genes can act as transcription factors (*e*.*g*., *malT, nadR*, and *iclR*) (1). We considered that the same change in metabolic phenotype could be achieved through a mutation in a regulator (altering expression levels) or direct mutation in a metabolic enzyme. To capture this idea, we generated adjusted mutation counts for each line, where a mutation in a regulator was propagated to a mutational count in each of its downstream targets, as identified by RegulonDB (29) (SI Appendix Section 9A). With these “regulatory adjusted” mutation counts, the outcomes of evolution in each line appear even more consistent, with an average correlation of 55% among the 3,807 non-regulatory genes (Figure 6B), suggesting a common direction of evolution. (While these increased correlations could be induced by the regulatory-adjustment process, we confirmed this effect was minimal by repeating the procedure with shuffled regulatory and regulated genes (SI Appendix Figure S14)).

The data are limited compared to what we can evaluate in simulation, but nevertheless the larger, second regime is sufficient to test for constant selective pressure over time. To do so, we turn to an empirical measure of the sensitivity of fitness to a mutation. In brief, we use the fact that mutations conferring a stronger advantage should rise in frequency in the population more quickly, allowing us to estimate the fitness sensitivity of each mutation by fitting a logistic curve to an individual mutation’s frequency trajectory (SI Appendix Figure S12). For a given window of time, we can then compute a proxy for the selective pressure on each gene through the accumulated fitness sensitivities of any mutation to that gene over the specified window. The fitness sensitivity to a direction of genotypic change can be calculated though a linear combination of these gene sensitivities.

Using this approach, we evaluated variance in fitness sensitivity across time and lines. We calculate the direction of change from mutation counts accumulated in the second regime (SI Appendix Section 9B). In short, we select the three lines with the largest average correlation and aggregated their regulatory-adjusted mutation counts in the second regime. Using the remaining three lines, we then evaluate fitness sensitivity to this direction over 5,000 generation windows (Figure 6C). Because the measure of selective pressure is imperfect and the data limited, for comparison we repeat the process using random permutations of the mutation count vector to estimate a null expectation (SI Appendix Section 9B). Computing the variance in fitness sensitivity for the true and permuted directions, we find the true direction of evolution has lower variance in fitness sensitivity than 99.5% of the randomly permuted vectors (Figure 6D).

In aggregate, the long-term evolution experiments show signs of being driven by EvCMs. Our results indicate common, reproducible directions of evolution emerge at higher levels of organization, and we find evidence for evolution along a constant selective pressure direction.

## Discussion

Our understanding of evolution is best in simple settings: a single gene controls a single phenotype that determines fitness. However, complex traits underlie fitness in every scale and domain of life. Understanding the evolution of such traits remains a central challenge in biology. The dominant approach of population genetics abstracts away the genotype-to-phenotype map, obfuscating or replacing complexity with marginal and linear effects. Here, we established a genotype-to-phenotype-to-fitness map underpinning a model complex trait (flux through a metabolic network), building on the successful and quantitatively tractable framework of flux balance analysis. Surprisingly, we find that explicit representation of this complexity in evolutionary simulation gives rise to simple and reproducible dynamics.

The EvCMs identified in our work bear a conceptual resemblance to the evolutionary sectors identified by Halabi, Rivoire, Leibler, and Ranganathan in protein families (30), where statistical analysis of sequence variation revealed groups of coevolving amino acids under collective rather than individual selection. Just as sectors identify collections of amino acids shaped by shared functional constraints (30, 31), our EvCMs identify collections of genes whose coordinated changes are favored by the structure of the metabolic genotype-phenotype map. The analogy is instructive: in both cases, biological constraints impose low-dimensional collective modes on an otherwise high-dimensional evolutionary space, suggesting that selection on collections — whether amino acids or genes — may be a general feature of the evolution of complex biological systems.

Though our setup is considerably simplified compared to living systems, we expect many of the lessons above will also generalize to more complex settings beyond metabolic networks and proteins. We identify collective modes, rather than individual genes, as the coherent and organizing objects of selection – a result we expect to hold in some general form when genes and phenotype contribute to fitness in a non-local, collective way. This could arise through developmental tradeoffs, common in many systems (*e*.*g*., growth versus defense in plants (32), investment in secondary sexual traits in beetles (33, 34), or warm versus cool adaptation in bacteria (35)), or through alternative instances of conservation constraints. In these cases, traits become coupled and evolutionary dynamics are not easily reduced to marginal impacts from individual genes. When collective mode-like dynamics are in play, we also anticipate that evolvability will be a fruitful lens to understand the favored direction among seemingly degenerate outcomes, and that evolutionary dynamics will transition through regimes based on evolvability and non-genetic constraints.

Taking a broader view, a defining feature of our framework is the centrality of phenotypic constraints in evolutionary analysis. Optimality has frequently been used as a lens to understand evolution but has little utility in our setting; without constraints, there is no well-defined optimum to the metabolic network. Subject to fixed constraints, phenotype and fitness may be optimized (with considerable degeneracy), but this teaches only about a relatively static picture of evolution. The constraints, therefore, dictate not only what is possible in relative evolutionary stasis, but also dictate the setting for longer-term dynamics. In light of this, analyzing the dual of evolution’s primal problem (*i*.*e*., shadow price vectors and not just phenotype vectors) becomes pivotal.

Taking a longer view, our work begins to clarify how evolution might act on the phenotypic constraints themselves. While we only considered scenarios where the constraints placed on phenotype were fixed, our work suggests that evolvability plays a key role in determining the EvCM. This observation suggests that evolution might act on constraints to increase evolvability. For example, might operons act to increase the evolvability of a collection of genes and be selected for? We believe our framework of constant selective pressure can help to answer this question and others regarding genetic structure.

Other aspects of this work may not generalize so readily. Most notably, despite degeneracy and randomness our simulation outcomes were remarkably consistent across replicate trials. This appears to be mirrored, at least to some extent, in the long-term evolution experiment with *E. coli*. What is absent from both settings is ecology or significant environmental feedback (though it has been observed that ecology emerged in the Lenski lines despite the experimental design to minimize ecological effects (36, 37)). Such feedback can in principle destabilize evolutionary trajectories (38, 39) so that “replaying the tape of life” would indeed produce new outcomes. Even if through some other mechanism, we do not in general expect outcomes to be as deterministic as demonstrated here. Further, a defining feature of collective modes in this setting – constant selective pressure – likely arises from the linearity of the system and might not manifest the same way in non-convex settings.

## Materials and Methods

Further discussion of methods is included in the SI Appendix.

### Evolutionary Simulations

As discussed in the main text, simulations were performed by mutating the genes of a metabolic network, using Equation 3 to calculate a new growth rate, and using Equation 1 to determine if the mutant goes extinct or fixes. Specifically, for a metabolic network described by *S, A, G*, and *β*, we start with a randomly selected value for the genes, **g**. The initial values of **g** were independently sampled from a normal distribution with a small variance. The small noise makes it easier to evolve initially as only a few reactions will be rate-limiting. Then, mutation is modeled by adding a random change Δ**g** to **g**. To sample Δ**g**, each gene was independently selected to be mutated with equal probability and, of the genes that were mutated, the effect of the mutation was sampled from a normal distribution with mean 0. One gene was mutated on average per attempted mutation. Next, the mutant phenotype **v**′ and fitness *f* ′ are again found using Equation 3. Then, *s* = *β*^*T*^ **v**′ − *β*^*T*^ **v**. Equation 1 was used to determine if the mutant fixes or goes extinct. This process is repeated for 10^6^ − 10^8^ steps to simulate an entire evolutionary trajectory.

### Dual of FBA and Selective Pressure

We used the dual of Equation 3 to estimate the selective pressure on a gene. The dual of the Equation 3 is

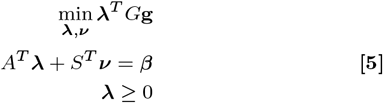

λ and *ν* are referred to as shadow prices and represent the sensitivity of the growth rate to changes in the constraints. Of importance to us, λ determines the sensitivity to a change in the constraints imposed by genes:

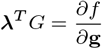

As mentioned, 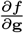 is the selective pressure on a gene. Then, we solve Equation 5 to obtain λ and use this to calculate the selective pressure on a gene. To calculate selective pressure on a *direction* (*e*.*g*., the EvCM), we weight the selective pressure of each gene by its contribution to the direction and sum: λ^*T*^ *G***z**. To calculate selective pressures over evolution, we resolve Equation 5 at each step. As **g** varies throughout evolution, so too will the solution to Equation 5 and value of λ.

## Data, Materials, and Software Availability

Code to perform simulations and example analysis to generate analysis for figure 2 are included on Github: https://github.com/algo-bio-lab/evolutionary-collective-modes.

## ACKNOWLEDGMENTS

We would like to thank members of the Cleary lab and Mehta group for useful discussions. AB was supported by NIH NIGMS 5T32GM145455-04. PM was supported by NIH NIGMS R35GM119461 and a Chan-Zuckerberg Investigator grant to PM.

## Supporting Information

### 1. Flux-balance analysis

#### A. Traditional FBA

Flux balance analysis (FBA) is a model to describe the metabolism of a cell (1). Instead of describing metabolism mechanistically (*e*.*g*., with kinetic parameters), FBA instead describes the feasible space of reaction fluxes and selects the ‘optimal’ set of reaction fluxes (2). In traditional FBA, the feasible space of reaction fluxes is primarily constrained by conservation of mass. The optimal set of reaction fluxes is the one that maximizes growth rate. This description can be written mathematically:

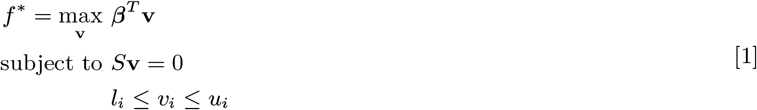

**v** is a vector of reaction fluxes for each reaction in the network. The growth rate is written as *β*^*T*^ **v**, so *β* is a vector that indicates how much each reaction contributes to the growth rate. Often, there is a “pseudoreaction” that represents the aggregate biomass metabolism (*e*.*g*., consume ATP, H_2_O, NAD+ and produce H+, ADP, and NADH to “grow”), so *β* is a vector of all 0s and one 1. The standard formulation of FBA seeks to maximize growth rate, so the problem maximizes *β*^*T*^ **v**. The optimal set of fluxes is **v**^∗^ and the optimal growth rate is *f* ^∗^ = *β*^*T*^ **v**^∗^.

The reaction fluxes must obey conservation of mass, and this requirement is encoded in *S***v** = 0. *S* is a stoichiometric matrix, where each column contains the stoichiometric coefficients of an individual reaction. Then, each row corresponds to a metabolite, and the size of matrix is the number of metabolites by the number of reactions: *m* × *r. S***v** = 0 imposes a steady-state condition on the fluxes; there is no net accumulation of any metabolites.

Beyond conservation of mass, additional bounds on individual reactions can be included, as represented by *l*_*i*_ ≤ *v*_*i*_ ≤ *u*_*i*_. Biologically, these bounds might enforce an irreversible reaction or encode an environmental limit on a exchange flux (*i*.*e*., a reaction that carries metabolites across the cell boundary) (1).

The traditional workflow of FBA is to solve Equation 1 to obtain the optimal set of reaction fluxes **v**^∗^ and growth rate *f* ^∗^. By varying features of Equation 1, metabolism under different conditions can be predicted. For example, knock-out and double knock-out viability has been explored by adding additional bounds (*i*.*e*., more bounds of the form *l*_*i*_ ≤ *v*_*i*_ ≤ *u*_*i*_) that force specific reaction fluxes to 0 (1).

#### B. FBA as a GP map

As described in the main text, we are interested in how genes evolve under a non-trivial genotypephenotype (GP) map. FBA (Equation 1) does not explicitly include genes. To use FBA as a quantitative description of a GP map, we augment traditional FBA by adding genes:

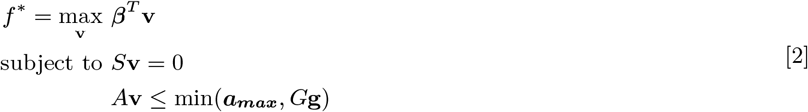

Like in traditional FBA, we maximize growth rate (*β*^*T*^ **v**) and require that the reactions must conserve mass at steady-state (*S***v** = 0). We incorporate the genes, represented by a vector **g**, into the bounds on reaction fluxes. Because metabolic fluxes are coupled across the metabolic network by conservation of mass, a gene cannot precisely set the reaction flux of a reaction it catalyzes. Instead, we assume that genes set limits on the reaction fluxes (*A***v** ≤ *G***g**). These limits could be set by the expression level of the gene or the inherent catalytic activity of the enzyme. Alternatively, ***a***_***max***_ sets a hard limit that can be used to incorporate environmental factors.

Each entry of **g** is associated with one gene and represents the maximum allowable flux of a gene: how fast a reaction *could* be run if the gene was fully expressed and reactants were abundant. Clearly, we are not modeling DNA sequences, but a more “downstream” feature of a gene. These maximum allowable fluxes then set “constraints” (*G***g**) on reaction fluxes. Put explicitly, there are three quantities of interest in Equation 2: reaction fluxes (**v**), constraints (*G***g**), and the maximum allowable fluxes of genes (**g**). In the main text, we primarily discuss reaction fluxes and maximum allowable fluxes, but the constraints are important as well.

While sometimes there is a one-to-one relationships between constraints and genes, often more than one gene is involved in catalyzing a reaction, leading to a potentially complicated relationship between each individual gene’s maximum allowable flux and the maximum allowable flux of a reaction. Including *A* and *G* allow for description of these complicated relationships, as discussed more below (Section 2).

### 2. Converting gene rules into constraints

In traditional FBA (Section 1A), the relationship between genes and reactions is described by a “gene rule” (3). A gene rule determines if a reaction can have a non-zero flux in a binary manner. In the simplest scenario, a gene rule is simply the absence or presence of a gene. If the gene is present, the flux can be non-zero; otherwise, the flux is forced to zero. However, more complicated versions can be constructed by generating Boolean expression of genes. For example, a gene rule might be “reaction flux can only be non-zero if gene A AND gene B are present.” We used these gene rules to parameterize our version of FBA (Section 1B), and, to do so, developed a process to translate the Boolean expression into linear inqualities. To describe this procedure we start with two examples and then discuss the general steps. Ultimately, this process motivates the inclusion of *A* and *G* in our verison of FBA.

#### A. Converting ORs

First, we focus on a scenario where genes rules are composed exclusively of ORs. For example the third step of glycolysis is the conversion of *β*-D-fructose 6-phosphate into *β*-D-fructose 1,6-bisphosphate (phosphofructosekinase reaction, PFK) (4). In *E. coli*, this reaction can be catalyzed by the proteins associated with two genes: *pfkA* and *pfkB* (5), which leads to the associated gene rule (3):

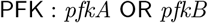

Broadly, we assume that protein abundance is the key factor that sets the maximum allowable flux for a reaction. So, we assume that the maximum allowable flux of each individual gene is additive, which leads to the following bound on the reaction:

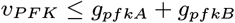

To again be clear, *g*_*pfkA*_ and *g*_*pfkB*_ are the maximum allowable fluxes of the gene, while *g*_*pfkA*_ + *g*_*pfkB*_ is the constraint. Incorporating the *G* matrix allows us to write the right-hand side.

#### B. Converting ANDs

A second example shows how to convert a gene rule composed of exclusively ANDs. The conversion of *α*-ketoglutarate to succinyl-CoA (*α*-ketoglutarate dehydrogenation, AKGDH) in the citric acid cycle of *E. coli* requires proteins from three genes: *lpd, sucB*, and *sucA* (6, 7). AKGDH has the following gene rule (3):

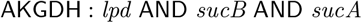

These proteins must join in stoichiometric proportion to form the reaction complex (8). If we again assume protein abundance is the primary factor in the total allowable flux associated with a reaction, then we expect that the least abundant protein would set the maximum allowable flux. Mathematically, this means the minimum of the maximum allowable fluxes of each gene sets constraint for the reaction:

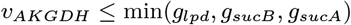

This minimum can be represented by three inequalities, which can be incorporated using the *A* matrix.

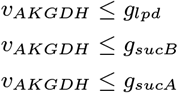

#### C. Converting combined ANDs and Ors

The above scenarios are examples where exclusively OR or AND relationships were used in the gene rules. In reality, some gene rules may have limits set by both operations. For example, the reaction carried out by cytochrome oxidase bd (CYTBD) has the following gene rule (3):

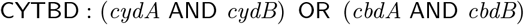

To consider more complicated scenarios like this one, we need to generalize the above process.

From the previous examples, we first have the following rules:

| Gene rule | Maximum Allowable Flux for Reaction R |
| --- | --- |
| R : A OR ... OR Z | $g_A + \dots + g_Z$ |
| R : A AND ... AND Z | $\min(g_A, \dots, g_Z)$ |

To use this mapping in more complicated scenarios, like for CYTBD, we needed to clarify how the OR and AND operations interact. With the gene rules, we assume they have their normal properties (*i*.*e*., they follow Boolean algebra). However, the maximum allowable fluxes do not follow the rules of Boolean algebra. Specifically, addition is not idempotent (while OR is):

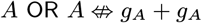

This fact also means that addition does not distribute over the minimum. As a result, all distribution of AND over OR must be undone before conversion from a gene rule to a maximum allowable flux. To see why, consider a hypothetical reaction with the following gene rule:

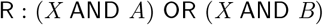

Directly converting from the gene rule would suggest that:

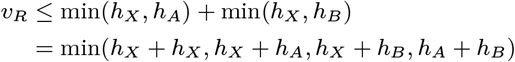

However, gene *X* cannot physically contribute double its maximum allowable flux, as seen in the *h*_*X*_ + *h*_*X*_ term. Instead, the distribution of AND over OR must be undone:

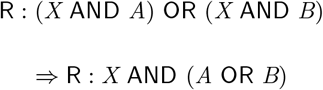

Then, the correct maximum allowable flux can be obtained:

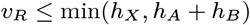

To undo the distribution of AND over OR, the gene rule can be converted to conjunctive normal form. Then, the maximum allowable flux can be obtained from the gene rule:

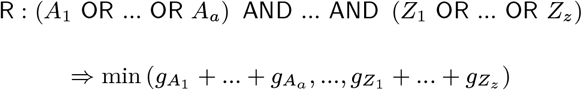

Summarizing, to turn a generic gene rule into a maximum allowable flux for a specific reaction, we first convert the gene rule into conjunctive normal form and simplify using the idempotent property of OR as necessary. Then, the maximum allowable flux of the genes associated by ORs are summed, and the constraint is set by the minimum of the sums.

Returning to the example with CYTBD, the conjunctive normal form of the gene rule shows that the maximum allowable flux is set by pairwise combinations of the individual genes:

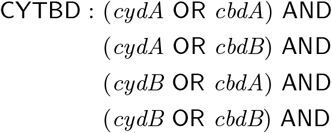

From the conjunctive normal form, the following maximum allowable flux for the reaction can be obtained:

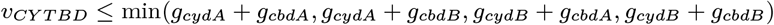

As discussed above, the minimum can be implemented by using four rows of *A*:

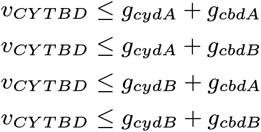

As can be seen, both *A* and *G* are necessary to implement these bounds.

### 3. Simulations of evolution of metabolic networks

Our goal is to understand how metabolic genes (**g**) evolve under the GP map described by our version of FBA (Section 1B, Equation 2). To this end, we simulated evolution of a metabolic network *in silico*, using Equation 2 as a a model of metabolism. As described in the main text, we performed sequential fixation simulations. After initializing the simulations, in each step we 1) generate a random mutation, 2) calculate the growth rate of this mutant, and 3) determine if it fixed with probability given by Kimura’s equation. These steps are repeated many times to simulate evolution, and are discussed in more detail below.

#### A. Simulation structure and hyperparameters

To model metabolism in the simulations, we explicitly separated upper and lower bounds, leading to a variation of Equation 2:

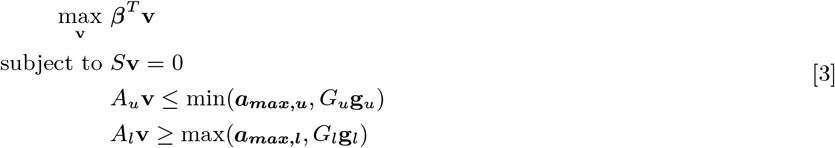

We partitioned the **g** vector into upper and lower genes, and assume that **g**_*u*_ ≥ 0 and **g**_*l*_ ≤ 0. This separation means that *G* is a 2 × 2 block matrix where the upper-left and lower-right sections correspond to *G*_*u*_ and *G*_*l*_, respectively, and the remaining entries are 0. Similarly, *A* can be divided into rows that corresponds to upper bounds (*A*_*u*_) and lower bounds (*A*_*l*_). ***a***_***max***_ is partitioned similarly.

The simulations we performed had various hyperparameters beyond the values of *S, A, G, β*, and ***a***_***max***_ needed to define a metabolic network. Here, we list key hyperparameters and their meanings:

1. *T* - the number of time steps in the simulation. *T >* 10^6^ for all simulations.
2. *N* - the population size. *N* = 10^7^ for all simulations (9).
3. *ss* - the simulation scale. This parameter sets the scale of the entire simulation. *ss* varied between 10-10,000. As further discussed below, increasing *ss* helped ensure evolution occurred.
4. *ms* - the mutation scale. This parameter sets the scale of mutation effects relative to the simulation scale. *ms* = 0.01-0.1 for the simulations.
5. *em* - the average number of genes mutated in a time step of the simulation. *em* = 1 across all simulations.

Hyperparameters were selected with the primary goal of observing any evolution, and, with knowledge of the existence of the EvCM, observing convergence. The length of simulation (*T*) that was necessary to see convergence to the EvCM appeared to scale with network size, so larger networks were run longer (*i*.*e*., compare the simulations of the *E. coli* central metabolism in Table S2 to the toy network). Similarly, scaling *ss* (in the context of *N* = 10^7^) primarily appeared to determine if evolution occurred at all, and was set to a value large enough to observe growth rate increasing for each simulation.

#### B. Initialization

To start the simulations, we generate initial maximum allowable fluxes (**g**_*u*,0_ and **g**_*l*,0_) and then solve Equation 3 to obtain the initial fluxes (**v**_0_). The maximum allowable fluxes were generated in two ways. For the toy networks (Section 4) and random metabolic networks (Section 7), we independently sampled from a normal distribution: **g**_***u***,**0**_ ∼ **𝒩 (*ss*1**,0.000025***ss***^**2**^***I***) and **g**_*l*,0_ ∼ 𝒩 (−*ss***1**,0.000025*ss*^2^*I)*. We then solve Equation 3 with the random bounds to obtain **v**_0_. For the simulations of the *E. coli* central metabolism (Section 8), we sampled **g**_*u*,0_ and **g**_*l*,0_ uniformly from [0, 1] and [−1, 0], respectively. Because the maximum fitness of the *E. coli* network is capped due to the limit on glucose intake, this approach was taken to observe as much of evolution as possible (*i*.*e*., observe evolution from a low fitness to its maximum, rather than only a fraction of the trajectory).

#### C. Mutation

At a simulation step *t*, mutants were generated by mutating the current genes **g**_*u,t*_ and **g**_*l,t*_. For both the upper and lower bounds, we sample a mutation Δ**g**_*u*_ and Δ**g**_*l*_. Broadly, Δ**g**_*u*_ and Δ**g**_*l*_ were generated to mutate one gene on average with a normally distributed effect. Specifically, we first independently select if each gene will be mutated with probability *p* = *em/*(|**g**_*u*_| + |**g**_*l*_|), where |*x*| is the dimensionality of *x* (*i*.*e*., *p* is 1/the total number of genes). As a result, *em* = 1 genes are mutated on average. Then, for each mutated gene, we sample a mutation effect from a normal distribution 𝒩 (0, *ms*^2^***ss***^2^). The mutations to upper and lower bounds Δ**g**_*u*_ and Δ**g**_*l*_ are then vectors of 0s for genes that were not selected to be mutated and normally distributed random effects for genes that are mutated. The mutant gene values are then 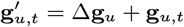 and 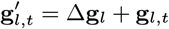.

Sometimes the above procedure can violate **g**_*u*_ ≥ 0 or **g**_*l*_ ≤ 0 (Section 3A). For example, if **g**_*u*_ = 0.5, but Δ**g**_*u*_ = −1, then 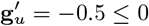. If this occurs, we force the values of 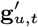 or 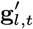 that violate **g**_*u*_ ≥ 0 or **g**_*l*_ ≤ 0 to 0. As most genes are increasing in magnitude due to evolution (*i*.*e*., see Figure S1A), this is often not a concern, and the above description of the mutation process is accurate.

#### D. Growth rate of mutant

With 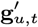 and 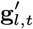, the growth rate of the mutant was calculated. Again, we use Equation 3 to model metabolism and determine the mutant growth rate. Explicitly,

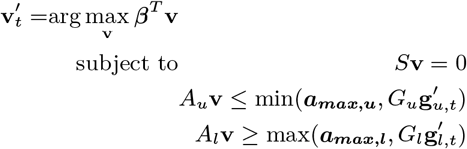

The fitness of the mutant is then 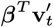.

#### E. Fixation of mutant

With the growth rate of the mutant calculated, Kimura’s probability of fixation was used to determine whether or not this new mutant will fix in the population:

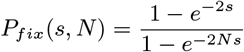

Kimura’s formula depends not only on the selection coefficient, 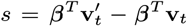, but also the total population size, *N*. Mutations that decrease fitness have a low chance of fixing in the population, while mutations that increase fitness have a larger chance. The population size also plays a role. Specifically, mutations have a larger chance to fix in smaller populations, regardless of selective advantage. As discussed above, *N* = 10^7^ for all simulations, so the probability of fixation is strongly influenced by the value of *s* and neutral mutations are rare.

The mutant fixes in the population with probability ***P***_***fix***_. If the mutant fixes, the genes and fluxes are updated: 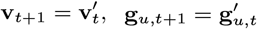 and 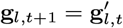. If the mutant goes extinct, the genes remain the same as the previous step, representing the mutation dying out in the population: **v**_*t*+1_ = **v**_*t*_, **g**_*u,t*+1_ = **g**_*u,t*_ and **g**_*l,t*+1_ = **g**_*l,t*_. This process is repeated for many time steps (*T* ≥ 10^6^) to model evolution.

### 4. Toy network and variants

As discussed in the main text, we performed simulations of a toy network to show the main results in a simple setting. A metabolic network is fully described by the parameters of our version of FBA (Section 1B) Here, we list *S, A, G, β*, and ***a***_***max***_ for the toy network and its variants. The hyperparameters are the same for all simulations of the toy network (Table S2).

#### A. Original toy network

Simulations of the original toy network were performed in Figure 2, Figure S1, and Figure S2. The parameters describing this network are:

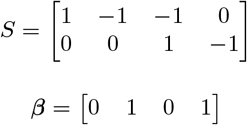

As mentioned above (Section 3A), both upper and lower constraints/genes are simulated. So, there are 8 constraints and 8 genes in this network. Specifically, *A* = [*A*_*u*_ − *A*_*l*_]^*T*^ = [*I* − *I*]^*T*^ and *G* = *I*. The first four rows of *A* and *G* correspond to the upper constraints, while the last four rows correspond to lower constraints. Because *G* = *I*, the first four columns correspond to upper genes, while the last four columns correspond to lower ones. ***a***_***max***_ = ∞ for all genes.

#### B. Toy network with flux limitations and regimes

To examine the role of limits on maximum flux imposed by the environment or gene expression, we used four variants of the toy network. The first two variants have the same *S, A, G*, and *β* as the original toy network (Section 4A), but ***a***_***max***_ ≤ ∞ for some genes to represent environmental (Figure 3A and B) and expression (Figure 3C and D) limitations:

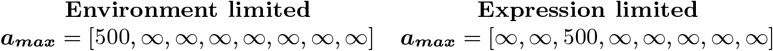

To show that more than two non-zero regimes are possible with one value of ***a***_***max***_ ≤ ∞, we constructed two additional variants of the toy network (Figure S3). Specifically, we found more than two regimes could arise if 1) different ‘pathways’ grow the cell different amounts (variation among *β*), 2) influx of nutrients is capped (***a***_***max***_ ≤ ∞), and 3) reactions cannot run in reverse (*a*_*max*_ = 0 for lower constraints). We list the values of parameters to achieve these conditions below:

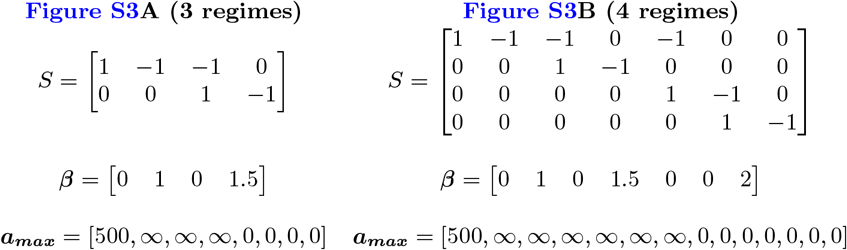

In both these networks, *A* = [*A*_*u*_ − *A*_*l*_]^*T*^ = [*I* − *I*]^*T*^ and *G* = *I*. Additionally, *a*_*max*_ for the lower constraints (the second half of ***a***_***max***_) is 0 for all reactions so that they cannot run in reverse. Forcing reactions to run forward is necessary, as running reactions in reverse could circumvent the nutrient limitation.

#### C. Toy network with variable number of independent genes

The toy network was changed to examine the role of a variable number of independent genes (Figure 3E). These variations of the original toy network (Section 4A) have the same value of *S, β*, and *A*. The values of *G* are:

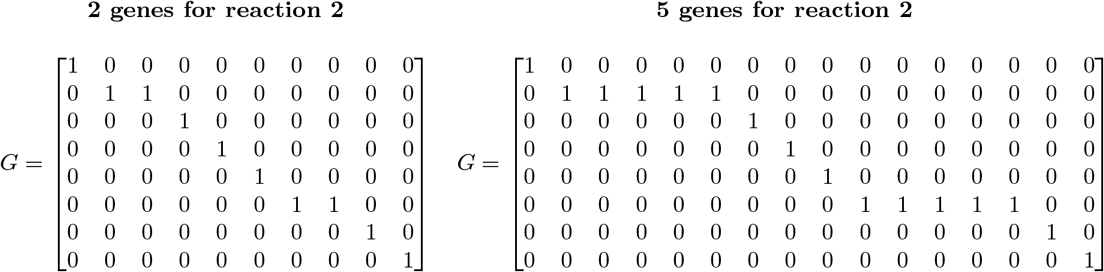

The extra genes were added for both the upper and lower bounds on reaction 2 (Section 3A), so there are 10 total genes and 16 total genes for 2 genes and 5 genes for reaction 2, respectively. Because reaction 2 is catalyzed by multiple independent genes, *G*≠ *I*. However, the number of constraints on each reaction has not changed, so *A* = [*I* − *I*]^*T*^, like in the toy network.

#### D. Toy network with variable number of dependent genes

The toy network was changed to examine the role of a variable number of dependent genes (Figure 3F). These variations of the original toy network (Section 4A) have the same value of *S* and *β*. The values of *A* and *G* are:

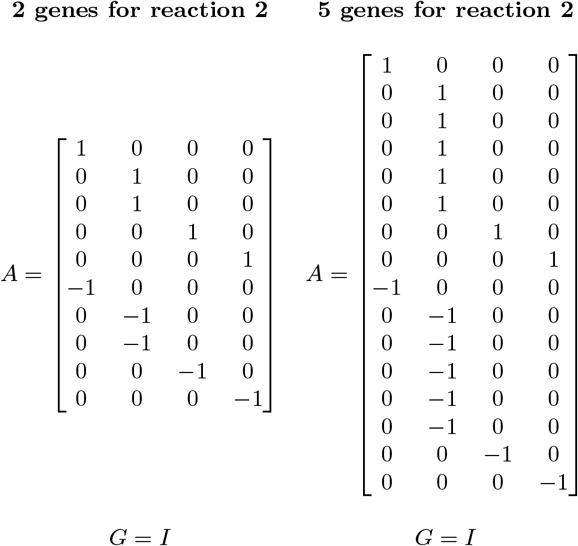

The extra genes were added for both the upper and lower bounds on reaction 2 (Section 3A), so there are 10 total genes and 16 total genes for 2 genes and 5 genes for reaction 2, respectively. Because each gene is only associated with one reaction and no reactions are catalyzed by multiple independent genes, *G* = *I*. However, the dependent relationship of the genes leads to extra constraints, so *A* [*I* − *I*]^*T*^, unlike the toy network; instead, there are 10 constraints and 16 constraints for 2 genes and 5 genes for reaction 2, respectively.

### 5. Selective pressure throughout evolution

As described in the main text, we observed the emergence of an evolutionary collective mode in our simulations. We wanted to understand how the EvCM arises, so we developed a mathematical framework to do so. In the context of our FBA-based description of metabolism (Equation 2), we want to understand how **g** changes as a result of evolution. Changes in **g** correspond to increasing or decrease selective pressure on genes, so we defined selective pressure on individual genes and combinations of genes, and then used duality to calculate the selective pressure.

#### A. Selective pressure on individual genes and directions

As discussed in the main text, we defined selective pressure (Δ*f*) on an individual gene *i* as the sensitivity of fitness to a change in the gene:

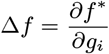

However, this framework does not allow us to consider the selective pressure on a combination of genes, like the EvCM. To calculate the selective pressure on a direction, Δ**g**, we assumed that each individual gene would contribute its selective pressure linearly:

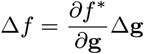

Here, both 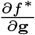 and Δ**g** are vectors. The definition of selective pressure for an individual gene can be recovered by setting Δ**g** to all 0s except for one 1 in the gene of interest. As mentioned, a combination of mutations can be encoded in Δ**g** (*i*.*e*., there are multiple non-zero entries). To ensure equal comparison between individual genes and directions, we force Δ**g** to have a 2-norm of 1 when we calculate selective pressure, so that we are only examining the selective pressure associated with a direction, not the number of mutations in the direction.

#### B. Calculation of selective pressure using duality

With selective pressure defined, we need to calculate 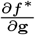, with *f* ^∗^(**g**) determined by Equation 2. Equation 2 is a linear program and we first focus on scenarios where *a*_***max***_ = ∞ so that it takes the traditional form of a linear program. Then, the concept of duality allows us to calculate 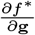. For every linear program, there is an associated linear program, called the dual (10). The dual of Equation 2 is:

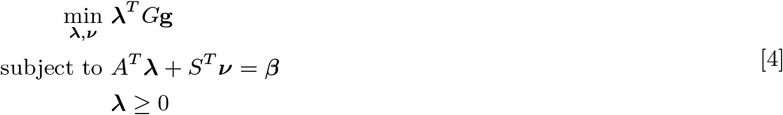

**λ** and ***ν*** are the dual variables that can be calculated given a network and current values of **g**. These variables capture the sensitivity of the optimal growth rate of Equation 2 to relaxations of the conservation of mass constraint or reaction bounds:

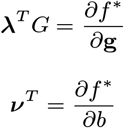

***b*** is the implicit parameter set to 0 in the conservation of mass constraint (*i*.*e*., ***S*v** = ***b***, with ***b*** = 0). **λ** measures the sensitivity to changes in constraints (*i*.*e*., *G***g**), so **λ**^***T***^ ***G*** measures the sensitivity to changes in genes. In biological terms, **λ**^***T***^ ***G*** is a vector that indicates, for each gene, how much growth rate will increase if the maximum allowable flux of a gene is increased. Similarly, ***ν*** describes the growth rate increase associated with a source of a metabolite appearing (being able to violate conservation of mass). We focus on the role of **λ**, as we are interested in the evolution of genes, but we hypothesize that ***ν*** may be relevant to understand the role of changing environments.

The definition of **λ** allows for calculation of selective pressure:

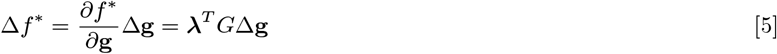

Equation 5 defines how we calculate the selective pressure on mutation Δ**g**.

##### B.1. Selective pressure with flux limits

The above discussion assumed that ***a***_***max***_ = ∞. However, when ***a***_***max***_ ≤ ∞, we still use the above definition of selective pressure, and we broadly motivate this as follows. A finite value for ***a***_***max***_ partitions the simulation into times when *G***g** ≤ ***a***_***max***_ and times when *G***g** ≥ ***a***_***max***_ (*i*.*e*., evolutionary regimes occur). When *G***g** ≤ ***a***_***max***_, ***a***_***max***_ does not affect the evolution, so the simulation essentially proceeds as before and the dual is the same, so the calculation of selective pressure is the same. When *G***g** ≥ ***a***_***max***_, the bound is fixed. Now, the dual is different, as it incorporates ***a***_***max***_ instead of the corresponding row of *G***g**, but it has the same general form. So, **λ**^***T***^ ***G*** still captures the sensitivity to mutations and can be used for the selective pressure on genes. However, now if a direction were to use the gene that cannot evolve due to ***a***_***max***_, the gene should instead be set to 0, as the constraint cannot evolve. Regardless, the definition of the selective pressure is the same.

#### C. Calculating selective pressure over partitioned upper and lower bounds

The above discussion of how to calculate selective pressure focused on a single *A* and *G*. As described above (Section 3), our simulations explicitly separated upper and lower bounds. Here, we show that the selective pressure can still be calculated using the dual and is similar to before. To see this is true, we first write the dual of Equation 3:

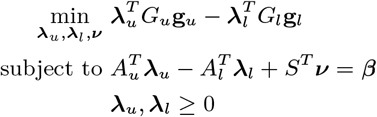

**λ**_***u***_ and **λ**_***l***_ describe the sensitivity of fitness to upper and lower genes:

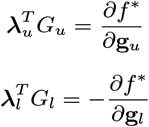

The negative sign is included with **λ**_***l***_, as increasing **g**_*l*_ is a tightening of a constraint, so ***f*** ^∗^ will generally decrease. Then, the selective pressure on a mutation with upper and lower components Δ**g**_*u*_ and Δ**g**_*l*_, respectively, is:

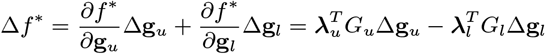

In the simulation, the values of **g**_*l*_ is negative, so the selective pressure Δ*f* ^∗^ is positive. However, in analysis, we often implicitly negate the lower bound components so that all elements are positive. Then, selective pressure is calculated as:

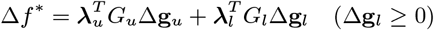

This definition of the selective pressure can also be seen as exploiting the block structure of *G* and assuming that **λ** can be partitioned as well.

### 6. Prediction of the EvCM

As described in the main text, we desired a way to predict the evolutionary collective mode (EvCM). In this section, we will write the EvCM as a single direction, **z**_*g*_, that is a concatenation of upper and negated lower gene values from simulation, as this is the most general scenario. After prediction, the individual elements can be re-attributed upper and lower genes if desired. We outline the next sections to give an overview. First, like before, we focus on scenarios where ***a***_***max***_ = ∞. Next, we find that many directions of evolution have constant selective pressure, so we posit that the EvCM is the one with maximal selective pressure. This assumption allows us to formulate an optimization problem to predict the EvCM, and we discuss approaches to solve it. Finally, we discuss scenarios were ***a***_***max***_ ≤ ∞ and derive a analytical result for the unique scenario where *A* = *I* and *G* = *I*.

#### A. Requirement of constant selective pressure

As described in the main text, our simulations suggested that constant selective pressure is a key constraint on the EvCM. We first asked how to mathematically describe this property and if this property is sufficient for predicting the EvCM.

The selective pressure on the EvCM is constant with respect to change in genes:

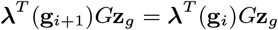

**g**_*i*_ are the gene values at simulation step *i*, and **λ** is written explicitly as a function of genes, as implied by Equation 4. To have constant selective pressure, this equality must hold at all steps up to the maximum step *T* :

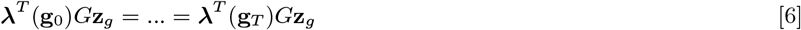

The ability of a single vector (**z**_*g*_) to satisfy the many equalities above is not apparent. *T* ∼ 10^6^, but the dimensionality of **z**_*g*_ is ≪ 10^3^ in our simulations, so conceivably, the ∼ *T* equalities should not be all satisfiable. This simple argument suggests that the relationship between **g** and **λ** is quite structured so that a single direction can indeed satisfy all the equalities. So, to determine which vectors have constant selective pressure, we need to better understand **λ**(**g**).

As discussed, **λ** as a function of **g** is determined by the dual of Equation 2 and its feasible region:

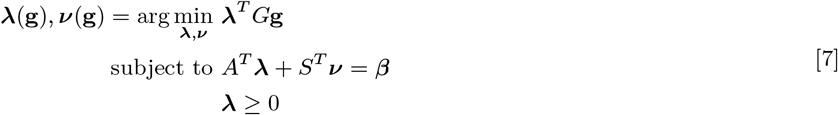

Because the optimization in Equation 7 is a linear program, solutions often occur at the vertices of the underlying feasible region, *P* :

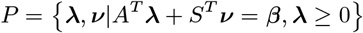

*P* is a polyhedron (11), and Equation 7 maps genes (**g**) to vertices of the polyhedron (**λ, *ν***). This suggests that the **λ** component of Equation 7 can be written in terms of the **λ** components of the *k* vertices of the polyhedron *P*, {**λ**_0_, **λ**_1_, …, **λ**_*k*_}:

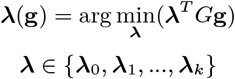

This reformulation of Equation 7 holds in most cases. However, **g** does not map to a single vertex if *G***g** is orthogonal to a face of dimension ≥ 1 of the polyhedron. If *G***g** is orthogonal to a face of dimension ≥ 1, then the objective value will be constant over that entire face, so any value of (**λ, *ν***) that is on the face is optimal. This scenario presents a potential challenge, as then a single value of **g** could map to many values of **λ**. However, evolution is a random process. The probability of a continuous, random vector being orthogonal to a face of the polyhedron is 0. So, a set of genes that result from evolution should almost always map to a vertex. This logic suggests viewing Equation 7 as a map from **g** to vertices is appropriate in our context.

Because there is a finite number of vertices on a polyhedron (11), Equation 6 can be reduced to equality over the vertices of the polyhedron:

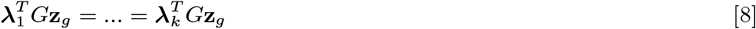

These *k* − 1 equalities can be restructured into a matrix equation:

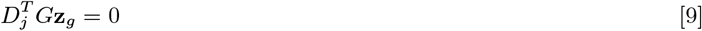

where

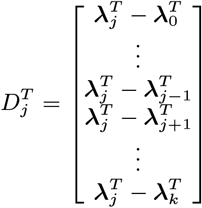

*j* indicates the shadow price that is the focal one in the definition of *D*_*j*_, and is arbitrary. Selecting any value of *j* describes the same solution space.

Equation 9 is a system of linear equations. So, three outcomes are possible: there are no solutions, there is exactly one solution, or there is infinite solutions. We observe directions with constant selective pressure, so seemingly there are indeed solutions. If there is exactly one solution, it is **z**_*g*_ = 0, which we also do not observe (genes accumulate mutations). Therefore, Equation 9 must have infinite solutions. This logic clarifies how constant selective pressure can be achieved at all simulation steps: **λ** only ever takes on a finite set of values (corresponding to vertices of the associated feasible-region polyhedron). Importantly, it also indicates that constant selective pressure is not a sufficient condition for predicting the collective mode; further assumptions must be made.

#### B. Requirement of maximal selective pressure

While we have clarified how constant selective pressure is achieved, the set of possible collective modes is still infinite. To identify the EvCM among the infinite solutions to Equation 9, we posited that a direction with constant *and maximal* selective pressure should be selected by evolution. Then, the EvCM is determined by an optimization problem:

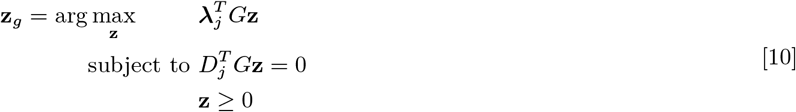

**z** ≥ 0 is included to account for physical feasibility. Equation 10 is unconstrained, but we can solve it under a norm to identify a direction:

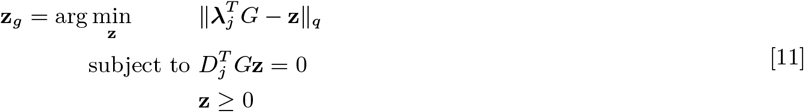

This problem is tractable once a value for the norm, *q* is selected. The challenge is then to determine **λ**_0_, …, **λ**_*k*_ so that we can solve Equation 11 to obtain **z**_*g*_. Once **λ**_0_, …, **λ**_*k*_ is determined, Equation 11 can be solve, and the EvCM can be predicted.

#### C. Determining λ_0_, …, λ_k_

As **λ**_0_, …, **λ**_*k*_ are the vertices of a polyhedron, one way to obtain **λ**_0_, …, **λ**_*k*_ is with a vertex enumeration algorithm, such as the double description algorithm (11). If this procedure is used for the toy network, the EvCM is correctly predicted. However, problems with this approach quickly arise. First, vertex enumeration scales poorly (12), so this approach is not useful once metabolic networks become large.

Additionally, even for smaller metabolic networks where **λ**_0_, …, **λ**_*k*_ can be enumerated, poor predictions are made for some metabolic networks. These inaccuracies occur seemingly because evolution does not visit every vertex of the polyhedron; selective pressure is only required over a subset of **λ**_0_, …, **λ**_*k*_. It is unclear how to identify which vertices are preferentially visited by evolution. Due to this challenge and the poor scaling, directly enumerating vertices to predict the EvCM is not useful.

An alternative way to obtain **λ**_0_, …, **λ**_*k*_ is to simply use the values of **λ** observed during an evolutionary simulation. At each step of evolution, **g**_*i*_ is used to calculate **λ**, and this process generates a list of **λ** that can be used to create 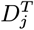. Then, Equation 11 can be solved. This strategy gives accurate results as shown in the main text with *q* = 2. To make a prediction at the level of constraints (*i*.*e*., predictions of *G***g**), we simply calculate **z**_*c*_ = *G***z**_*g*_. Again, this strategy gives accurate results with *q* = 2, as shown in Figure S13.

##### C.1. Chain sampling

While using the values of **λ** from simulation achieved accurate results, *a priori* prediction from the metabolic network is desirable. To address this challenge, we developed a sampling-based approach to identify **λ**. We use the following sampling scheme. First, we randomly sample a value of genes **g**^0^ (superscript denotes sampling steps). Then, solve Equation 4 to obtain **λ**^0^. Then, **g**^1^ = *ϵG*^*T*^ **λ**^0^ + **g**^0^, where *ϵ* is a small constant. Now, solve Equation 4 again to obtain **λ**^1^. Repeat this process, using **λ**^*i*+1^ = *ϵG*^*T*^ **λ**^*i*^ + **g**^*i*^. This strategy is motivated by the fact that **λ**^*T*^ *G* = *∂f* ^∗^*/∂***g**, so adding *G*^*T*^ **λ**^*i*^ approximates a fitness-increasing step.

After *C* samples, we take the unique values of {**λ**^0^, …, **λ**^*C*^}. To avoid errors with numerical precision, we rounded the values of **λ** to 5 decimal places before finding the unique values. We sort these unique values by the number of occurrences, and take the set **λ**_0_, …, {**λ**_*k*_} as all unique values of **λ** that occur with frequency greater than *α*. We used *α* = 0.001, *C* = 5000, and *ϵ* = 0.01. A small value for *α* was used to prioritize only removing infrequent **λ**s, while *ϵ* was selected to be small so that **g**^*i*^ would smoothly transition between regions corresponding to different **λ**s. While less accurate than using **λ** from simulation, this approach still gives accurate results with *q* = 2, as shown in Figure S13.

#### D. Prediction of EvCM with fixed genes

The above discussion assumes that no genes with fixed maximum allowable flux are present (*i*.*e*., ***a***_***max***_ = ∞ for all genes). However, we also make predictions when some genes do have finite values for ***a***_***max***_. In these scenarios, we assume that the maximum allowable flux of the gene was much larger than ***a***_***max***_ so the gene does not evolve. Then, we augment Equation 11 with an additional constraint that forces genes with finite ***a***_***max***_ to be 0 in the EvCM:

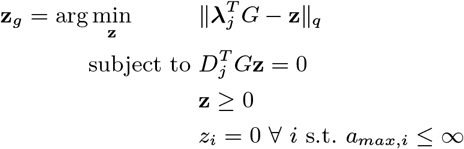

The addition of this constraint is motivated by the fact that the EvCM is a direction and when a gene cannot evolve, evolution cannot move in that direction. Otherwise, the logic is the same: select the direction with constant and maximal selective pressure. Figure S3G and Figure S9 show predictions for the toy networks with regimes and the central metabolism of *E. coli* (*i*.*e*., scenarios when ***a***_***max***_ ≤ ∞) using **λ** from simulation.

#### E. Prediction of EvCM

**when** *A* = *I* **and** *G* = *I*. As suggested by the above discussion, developing an analytical expression for the EvCM is challenging. However, in the specific scenario where *A* = *I* and *G* = *I*, an analytical prediction can in fact be made and provides some intuition about what shapes the collective mode. Additionally, in this scenario, as there is a one-to-one relationship between genes, constraints, and fluxes, we may be able to interpret the prediction as a *flux*-level prediction, **z**_*f*_.

As before, we begin by analyzing the dual (Equation 4). If *A* = *I* and *G* = *I*, the dual problem becomes:

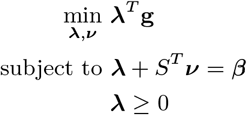

The first constraint (**λ** + *S*^*T*^ ***ν*** = *β*) means that **λ** = *β* − *S*^*T*^ ***ν***, so selective pressure can be defined in terms of ***ν***:

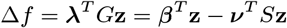

For a direction to have constant selective pressure, we then require, similar to before, for all pairs of ***ν***_*i*_ and ***ν***_*j*_, that:

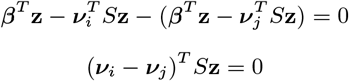

Like before, by combining the various equalities, we can convert to matrix form:

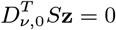

With these simplifications, Equation 10 can be written as:

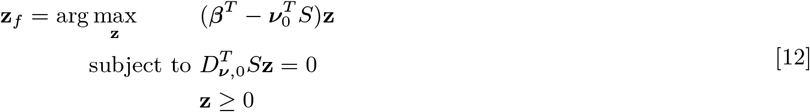

Currently, Equation 12 is no more intuitive than the original formulation of predicting the collective mode (Equation 10). However, we argue this problem can be further simplified. Based on Equation 12, one way to have constant selective pressure is to require *S***z** = 0. First, we note that this requirement is nominally not necessary, as **z**_*f*_ can be interpreted as genes, not fluxes.

Second, we note that vectors in the null space of 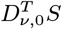 would also have selective pressure. However, we argue that 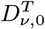 is full rank and *S***z** = 0 is sufficient for constant selective pressure. We make this argument based on the dimensionality of 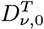. The width of this matrix is the number of metabolites, *m*. The height of this matrix is related to the number of specific values of ***ν***. Each value of ***ν*** indicates when a metabolite is limiting fitness. This description suggests that there will be at least *m* value of ***ν*** corresponding to a unique metabolite limiting fitness. More values will appear as combinations of metabolites begin to limit fitness. Therefore, 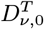 is a “tall-and-skinny” matrix and full rank. Importantly, 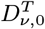 is a different matrix than 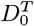, which is clearly not full rank based on our previous discussion.

If we enforce *S***z** = 0, Equation 12 is simplified:

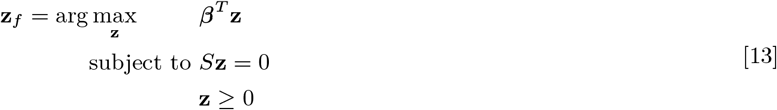

As before, we can solve Equation 13 under a constrained norm:

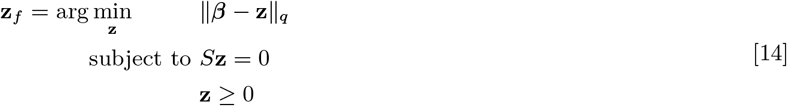

Because of the constraint **z** ≥ 0, Equation 14 still does not have an analytical solution. However, we argue that **z** ≥ 0 can be removed. We first note that discussion of the EvCM only makes sense if the original linear program (Equation 2) is bounded. Otherwise, infinite fitness would be achieved immediately. Generically, finite fitness requires that each reaction has both an upper and lower bound. In Equation 2, upper bounds correspond to when rows of *A* have positive entries, and lower bounds correspond to when rows of *A* have negative entries. By enforcing *A* = *I*, we implicitly assumed that only upper-bound constraints are necessary for Equation 2 to be finite. Said another way, the evolutionarily “preferred” direction of each reaction is the forward direction with positive flux. Returning to Equation 14, this discussion suggests that the constraint **z** ≥ 0 is redundant; the EvCM, **z**_*f*_, will have positive entries because the preferred direction for each reaction is forward.

If the constraint that **z** ≥ 0 is removed, the solution to this problem with *q* = 2 is the projection of *β* into Null *S*:

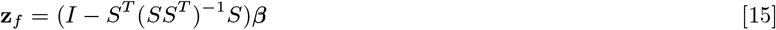

This result matches intuitive expectations. If a single direction of evolution were to exist at the flux level, it makes some sense that it should follow *β* and obey conservation of mass.

The above discussion suggests that this expression for **z**_*f*_ should only apply in scenarios where **z**_*f*_ ≥ 0. However, if we use this method when *G* = *I* and each reaction, except the biomass reaction, only has an upper and lower bound constraint, we achieve correlations between simulation and prediction *>*0.94 at the level of fluxes (Figure S13), even when though *S* is random and preferred directions of fluxes could be positive or negative (*i*.*e*., we have no guarantee **z**_*f*_ ≥ 0). We motivate this result with two arguments. First, if an entry of **z**_*f*_ is negative, the corresponding column of *S* can be negated, which will negate the entry of **z**_*f*_, so that **z**_*f*_ ≥ 0. Second, because evolution seemingly has a single preferred direction (*i*.*e*., the EvCM), only one of an upper or lower bound are active over evolutionary time scales. This feature suggests that one of the constraints is redundant to the problem, and the corresponding row in *A* could be removed without changing the results of evolution at the flux level.

The above discussion motivates that **z**_*f*_ is the projection of *β* into Null *S* when *G* = *I* and constraints on reactions are rows of the identity matrix. We use this solution for comparison to scenarios where extra subunits (*i*.*e*., *A* ≠ *I*) and extra genes (*i*.*e*., *G* ≠ *I*) are present in our discussion of evolvability.

Interestingly, this approach also works well when *A* ≠ *I* or *G* ≠ *I* (Figure S13) for predictions at the flux level. However, a majority of simulations analyzed (199/250) had more accurate result when collecting **λ** from simulation and using Equation 11. This result suggests that the framework of constant and maximal selective pressure is the correct one for predicting the collective mode, especially when *A* ≠ *I* or *G* ≠ *I*.

### 7. Random metabolic networks

#### A. Generating random metabolic networks

In our setting, a metabolic network is defined by *β, S, A*, and *G*, and we start here with the scenario ***a***_***max***_ = ∞. The size of each parameter is determined by four values:

1. *m* - the number of metabolites
2. *r* - the number of reactions
3. *c* - the number of constraints (2*c* constraints across upper and lower bounds)
4. *h* - the number of genes (2*h* genes across upper and lower bounds)

##### A.1. S. S is a m × r matrix

Generating *S* requires 5 parameters:

1. *n*_*import*_ - the number of reactions that will be import exchange reactions.
2. *n*_*export*_ - the number of reactions that will be export exchange reactions.
3. *z*_*max*_ - the maximum number of metabolites participating in one reaction.
4. ***ν***_*max*_ - the maximum stoichiometric coefficient of a metabolite.
5. *n*_*bio*_ - the number of metabolites participating in the biomass reaction.

Import reactions are generated by randomly selecting *n*_*import*_ metabolites from the total *m* metabolites and assigning a stoichiometric coefficient of 1. Export reactions are assigned analogously with *n*_*export*_ metabolites and a coefficient of − 1. Internal (non-exchange) reactions are generated to ensure that each metabolite is used at least once and every reaction both consumes and produces at least one metabolite. There will be *r* − 1 − *n*_*import*_ − *n*_*export*_ internal reactions. The number of metabolites participating is selected uniformly from [2, *z*_*max*_]. Stoichiometric coefficients for each metabolite are selected uniformly from [ − ***ν***_*max*_, − 1] ∪ [1, ***ν***_*max*_].

The biomass reaction is a special reaction. *n*_*bio*_ metabolites are selected to participate in it, and the stoichiometric coefficients are randomly selected from a standard normal distribution. The coefficients are then transformed to have a mean of 0, and the values are rounded to three decimals.

As mentioned in the main text, a randomly generated metabolic network might not evolve. This occurs if *β* is orthogonal to Null *S*. Then, the growth rate would be forced to 0, as *S***v** = 0, so **v** ∈ Null *S*, and *β*^*T*^ **v** = 0. We want to observe evolution, so if *β* is orthogonal to Null *S*, we repeat this procedure until we find a *S* that can evolve.

##### A.2. *β. β* is a vector with *r* entries

We use “biomass” reactions for all randomly generated metabolic networks, so β is a vector of 0s with one 1 corresponding to the biomass reaction.

##### A.3.A. *A* is a *c* × *r* matrix

As mentioned above, we are primarily interested in scenarios where reactions do not directly compete with each other. As a result, rows of *A* have one non-zero entry. So, we generate *A* by randomly selecting a reaction for each constraint. We do not impose any scale on the constraints, so all values of *A* are either 0 or 1: the constraint either does (1) or does not (0) limit the reaction. We do not impose any constraints on the biomass reaction (*i*.*e*., the corresponding column of *A* is all zeros), but it is indirectly constrained due to conservation of mass and constraints on the other reactions. We used this procedure to generate *A*_*u*_ and *A*_*l*_ separately.

##### A.4.G. *G* is a *c* × *h* matrix

There is one parameter in generating *G*: *p*, the minimum average number of genes per constraint. Genes are randomly assigned to constraints, and then additional relationships between genes and constraints are added until the average number of genes per constraint in *G* exceeds *p*. As when generating *A*, we do not impose a scale, so all values of *G* are either 0 or 1: the gene either does (1) or does not (0) control the constraint. We used this procedure to generate *G*_*u*_ and *G*_*l*_ separately.

#### B. Simulation

The simulations were performed as described in Section 3. In total, 400 random metabolic networks of varying size were generated and simulated. Table S3 lists the values of parameters discussed in Section 7A used to generate the random metabolic networks. The hyperparameters (Section 3A) used for the simulations of random metabolic networks are shown in Table S2.

#### C. Evolvability analysis

As discussed in the main text, we analyzed the random metabolic networks to understand how evolvability shapes the collective mode. To perform this analysis, we compared the EvCM of a random network to the EvCM predicted if *A* = *I* and *G* = *I* (Section 6E Equation 15). This process is conceptually similar to what was done in the main text with the toy network. The logic is that *A* ≠ *I* or *G* ≠ *I* should change the evolvability of reactions compared to when *A* = *I* and *G* = *I*. This change in evolvability should manifest in changes in the EvCM. By examining which features of *A* and *G* correlate with deviations, we can start to understand how evolvability is shaping the EvCM.

To analyze the role of evolvability, we performed specific simulations of random metabolic networks (more details are discussed below). Then, we calculated the normalized change in flux throughout the simulation (*i*.*e*., the EvCM):

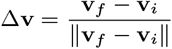

We also calculated the normalized prediction using Equation 15:

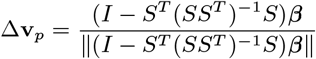

As mentioned, we attribute the difference between the EvCM and the prediction, Δ**v** − Δ**v**_*p*_, to evolvability. Below, we discuss how we constructed the random simulations to isolate specific types of evolvability.

##### C.1. Role of multiple dependent genes

As discussed in the main text, we hypothesized that reactions that require multiple dependent genes should be less evolvable and, as a result, have a smaller role in the EvCM. To test this, we performed simulations of random metabolic network where some reactions have gene rules where independent genes are joined by AND (*e*.*g*., *A* AND *B* AND *C*). These networks were created by simply requiring that there be more constraints than reactions while *G* = *I*. Specifically, we examined metabolic networks with *m* = 25 metabolites, *r* = 50 reactions, *c* = 75 constraints, and *h* = 75 genes. As mentioned above, *A*_*u*_ and *A*_*l*_ were generated separately. We performed simulations of 50 random metabolic networks (Table S3).

For each simulation, we calculated Δ**v** − Δ**v**_*p*_, as discussed above. Then, for each reaction, we determined if the upper or lower bound was active by examining the sign of Δ**v** (*i*.*e*., Δ**v** *>* 0 implies the upper bound and vice versa). For each reaction, we examined the active bound and counted the number of dependent genes (the number of constraints on that reaction). This process generates the data shown in Figure S8A. As discussed in the main text, we concluded that requiring more dependent genes lowers evolvability and leads to a lower role in the collective mode.

##### C.2. Role of multiple independent genes

As discussed in the main text, we hypothesized that a reaction that can be catalyzed by multiple independent genes should be more evolvable and, as a result, have a larger role in the EvCM. To test this, we performed simulations of random metabolic network where some reactions have gene rules where independent genes are joined by OR (*e*.*g*., *A* OR *B* OR *C*). These networks were created by simply requiring that there be more genes than reactions while *A* = *I*. We used *p* = 1.5 when generating *G*. Specifically, we examined metabolic networks with *m* = 25 metabolites, *r* = 50 reactions, *c* = 49 constraints (the biomass reaction is unconstrained), and *h* = 75 genes. As mentioned above, *G*_*u*_ and *G*_*l*_ were generated separately. We performed simulations of 50 random metabolic networks (Table S3).

For each simulation, we calculated Δ**v** − Δ**v**_*p*_, as discussed above. Then, for each reaction, we determined if the upper or lower bound was active by examining the sign of Δ**v** (*i*.*e*., Δ**v** *>* 0 implies the upper bound and vice versa). For each reaction, we examined the active bound and counted the number of genes that can catalyze the reaction (the sum of the corresponding row of *G*). This process generates the data shown in Figure S8B. As discussed in the main text, we concluded that using more independent genes increases evolvability and leads to a larger role in the collective mode.

### 8. Central metabolism of *E. coli*

#### A. Simulation

As described in the main text, we performed simulations of the central metabolism of *E. coli*. The simulations of the central metabolism of *E. coli* were carried out as described above. The “e_coli_core” model from BiGG was used (3). We generated *A* and *G* using the procedures for gene rules described above (Section 2). Simulations were performed at a simulation scale (*ss*) of 10,000 to increase the likelihood of beneficial mutations fixing and shorten the required number of simulation steps. The other hyperparameters used are included in Table S2.

We made some slight changes to the “e_coli_core” model. First, the “e_coli_core” model includes an “ATP maintenance” reaction with a fixed bound to represent an excess amount of ATP that must be generated that does not go towards biomass (1). As this limits the lower bound of fitness, we set this fixed bound to 0 (*a*_*max*_ = 0), and the evolution of the ATP maintenance reaction was not simulated (*i*.*e*., the ATP maintenance flux was always 0). Second, the central metabolism of *E. coli* contains exchange reactions and irreversible reactions. Both of these types of reaction share the common feature that one bound is fixed to 0 (*i*.*e*., the reactions run exclusively in one direction). To implement these reactions, constraints were incorporated that did not have an associated gene (so the flux is bounded by 0). Third, the exchange rate of glucose has a fixed upper bound of 10 (*i*.*e*., the environment has a limited amount of glucose). To match the simulation scale, the value of *a*_*max*_ for the glucose exchange was scaled as well and set to 10,000.

#### B. Analysis

Broadly, the simulations of the *E. coli* central metabolism were analyzed like the random metabolic networks. However, two features made analysis difficult: a period before evolution began and regimes.

The *E. coli* simulations all had a period where no evolution occurred before fitness started increasing meaningfully. We do not believe this period is meaningful and do believe that it may be an artifact of how simulations are initialized. As a result, we estimated the start of evolution for each replicate and independently aligned the simulations so that the start of evolution was time 0 in all replicates. Table S1 shows when evolution was estimated to start for each simulation.

The simulations of the *E. coli* central metabolism also contain regimes. The presence of regimes complicates the analysis of the simulation, as each regime has a EvCM. As a result, we analyzed the simulations within regimes, and the boundaries of the regimes had to be estimated. As mentioned in the main text, we looked for large changes in the direction of evolution as evidence of a transition between regimes. Table S1 shows where each regime was estimated to end. Because the simulations fall out of sync throughout evolution, the regime start times vary. Due to the imprecision in estimating where regimes start and end, values of **λ** were only taken from the middle 60% of estimated regimes when predicting the EvCM in Figure S9.

### 9. Lenski lines

As discussed in the main text, we analyzed the Lenski lines for evidence of evolution along an EvCM. Metagenomic sequencing has been performed every 500 generations for all lines up to 60,000 generations (9). This sequencing identified various mutations that occurred and their frequency across time, and we used this data to look for the impact of an EvCM.

Of the 12 populations in the Lenski lines, six have developed a hypermutator phenotype. As it is difficult to associate mutations to fitness benefits in the hypermutator lines, we did not analyze these lines. Instead, we focused on the normal mutators, where mutations occur less often and can be more clearly associated with driving changes in mutation frequency.

To obtain the data for analysis, we downloaded the processed data from (9) and modified their code to extract mutation frequency trajectories. We also used the modified code to generate the fitness data shown in Figure 6A. ChatGPT-4o was used to change code written for python 2 into code written for python 3.

#### A. Regulatory-adjusted counts

We first analyzed the Lenski lines by looking at correlations of mutation counts across lines. We saw little correlations across normal mutator lines (Figure S11). However, many regulatory genes are mutated in the Lenski lines, so we asked if correlations would appear if we examined only the downstream genes. To do this, we needed to collate the regulatory genes and their downstream targets. We used RegulonDB to collect this data (13). RegulonDB has regulatory relationships organized into “regulons.” So, for each regulon, we marked the genes that regulate it as “regulatory” and the downstream regulated genes as “regulated.” This data was assembled into a matrix with regulatory genes along the columns and regulated genes along the rows. Then, an entry of the matrix was set to 1 if there was a regulon where the regulatory gene (column) regulated the regulated gene (row). The regulatory data was collected from RegulonDB on May 22, 2025.

Using the data from RegulonDB, we recounted the mutations. If a mutation occurred in a regulatory gene, we counted it as a mutation in each of its downstream targets. As the effects of regulatory genes are included in this process, we did not also include regulatory genes in the regulatory-adjusted counts. When we examined the correlations across lines between regulatory-adjusted counts, we saw increased correlations compared to the raw mutation counts, as discussed in the main text and in Figure 6B.

To confirm that these increased correlations did not result exclusively from the regulatory-adjustment process, we generated a null distribution for comparison. Specifically, we mapped each gene (regulator or regulated) to another gene and swapped their role in the regulatory relationships. For example, if gene *A* regulated genes *B* and *C*, and gene *D* regulated gene *B*, one possible map could be {*A* → *D, B* → *A, C* → *B, D* → *C*}. Then, the shuffled regulatory relationship would be gene *D* regulates genes *A* and *B*, and gene *C* regulates gene *A*. Then, we recalculated the regulatory-adjusted mutation counts using this new mapping between regulatory and regulated genes. Finally, we calculated the correlation between lines. If the increased correlations were induced by the regulatory adjustment, we expected that the correlations with randomly swapped regulators would be increased as well. Instead, we found that many of the true regulatory-adjusted correlations between lines were in the top 10% of the distributions generated by random swapping (Figure S14).

#### B. Selective pressure

We next looked for evidence of constant selective pressure in the Lenski lines across both time and lines. Calculating a selective pressure requires a direction of evolution of genes (**w**) and a sensitivity of fitness to mutations (σ). Both these variables are vectors. Each element of **w** is the number of mutations in a gene, and each element of σ represents the fitness gain in the current state associated with mutating a gene. The selective pressure on the direction can be estimated as σ^*T*^ **w**. This calculation is similar in spirit to how we calculated selective pressure in simulations (*i*.*e*., **λ**^*T*^ *G***g**), but we use σ and **w** to emphasize that these calculations do not rely on the FBA-based description of metabolism from before.

As further described below, to calculate the selective pressure, we estimated one direction of evolution, **w**, using three lines (lines p1, p5, and p2). Then, we estimated values of σ over time. Specifically, we binned the data into 5,000 generation windows and calculated a value of σ within each window and for each line that was not used to estimate **w** (lines m5, m6, and p4).

To determine which mutations should go in which windows, we estimated when mutations appeared. For each mutation frequency trajectory in the data set, we set the appearance time as the earliest generation at which the mutation passed a frequency of 0.1. Additionally, as discussed in the main text, we needed to determine if any regimes were present. Clearly, the fitness increases non-linearly. We assumed that there were two regimes and focused on the second regime. The slope of the fitness appears to change at ∼ 2, 500 generations. To avoid any overlapping region, we performed analysis using data from ≥ 10, 000 generations.

##### B.1. Estimating w

**w** was the easier of the two variables to estimate. Using the lines saved for **w** (lines p1, p5, and p2), we calculated the regulatory-adjusted counts for each line in the second regime. We estimated **w** as the average mutation count for each gene over the three lines.

##### B.2. Estimating σ

To estimate σ, we first estimated the selective advantage associated with each mutation that occurred in the lines saved for σ (lines m5, m6, and p4). Specifically, we fit the frequency trajectory *f* of each mutation to a linearized logistic curve, where the slope is the selective advantage (Figure S12) (14):

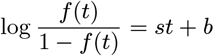

To improve the estimate accuracy, we took two steps to clean the data. First, we applied a median filter to the data to reduce small fluctuations in the frequency of a mutation. A window size of 3 was used. Second, as the logistic function is monotonically increasing, we only performed regression up to the time when the maximum frequency of the mutation occurs. Specifically, once the maximum frequency was recorded, any frequencies at later times were not included in the regression.

After estimating the selective advantage of each mutation, each mutation was assigned to a window based on its appearance time. Then, within a window, σ was generated by aggregating selective advantages of mutations in the same gene. For regulatory genes, the selective advantage was partitioned equally across its downstream targets. So, for each gene, the total selective advantage is the sum of the selective advantage for each mutation in that gene and the selective advantage contributed from its regulatory genes. This process was repeated across lines. As a result, there was 30 estimates of σ across time (10×) and lines (3×).

##### B.3. Estimating selective pressure

With one estimate of **w** and 30 estimates of σ, 30 estimates of the selective pressure on **w** were obtained by calculating **w**^*T*^ σ for each value of σ. This process generates the data shown in the EvCM portion of Figure 6C. The selective pressure on individual genes was simply the corresponding element of σ for that time and line. As **w** was not norm 1, the element of σ was scaled by the norm of **w**.

##### B.4. Evidence of constant selective pressure

Having calculated the selective pressure on the estimated EvCM of the Lenski lines, we looked for evidence of constant selective pressure. Using the previous estimates of the selective pressure, we calculated the coefficient of variation of the selective pressure across time and lines (*i*.*e*., **w**^*T*^ σ_1_, **w**^*T*^ σ_2_…**w**^*T*^ σ_30_). We expect this value to be small.

For comparison, we generated a null distribution. Specifically, we permuted the value of **w** by swapping elements, and then recalculated the selective pressure using the previously calculated values of σ. Then, we calculated a coefficient of variation among these selective pressures. Repeating this process many times created a null distribution. The comparison between the null distribution and the observed value is shown in the main text in Figure 6D.

### 10. General analysis of metabolic networks

The discussion above has primarily focused on analysis performed within the selective pressure framework (Section 5). Here, we describe other types of analysis we performed.

#### A. Identification of constant selective advantage by linear regression

The biomass of the metabolic networks appeared to increase linearly across time. To confirm this observation, we performed linear regression. Specifically, we fit the biomass *f* as a function of simulation steps *t*:

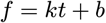

We calculated *R*^2^ for each fit to quantify if fitness increased linearly.

#### B. Estimation of EvCM from simulation

The EvCM of a metabolic network was calculated as the change in flux or maximum allowable flux over the entire simulation, unless otherwise indicated. Specifically, we calculated:

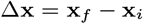

**x**_*f*_ is the last value in the simulation, while **x**_*i*_ is the first value. This calculation is performed with fluxes for reactions or maximum allowable fluxes with genes. Often, the normalized direction is used in downstream analysis, which is calculated as: Δ**x***/*∥ Δ**x** ∥_2_. When normalizing a change in maximum allowable fluxes, the upper and lower genes are concatenated and the norm of this vector is set to 1.

#### C. Identification of constant direction

To identify if a metabolic network evolved along a constant direction, we examined the change in fluxes/maximum allowable fluxes across a short time period. If this direction is constant across time, then the network is evolving along a constant direction. How constant the direction appears is impacted by the length of the time period examined, so we list the time periods used below:

1. Toy network (Figure 2C, Figure S1B) - 50,000 attempted mutations.
2. Toy network with regimes (Figure 3B) - 2,000 attempted mutations.
3. Toy network with regimes (Figure 3D) - 5,000 attempted mutations.
4. Toy network with regimes (Figure S3D) - 5,000 attempted mutations.
5. Toy network with regimes (Figure S3F) - 5,000 attempted mutations.
6. E. coli regime 1 (Figure S10) - 1,000,000 attempted mutations.
7. E. coli regime 2 - 700,000 attempted mutations.
8. E. coli regime 3 - 700,000 attempted mutations.
9. E. coli regime 4 - 20,000,000 attempted mutations.
10. E. coli regime 5 - 2,000,000 attempted mutations.
11. E. coli regime 6 - 10,000,000 attempted mutations.
12. Random replicates (Figure S7) - 500,000 attempted mutations.

In some figures, we compare the change in fluxes/maximum allowable fluxes (Δ**x**) to the estimated EvCM (**x**_*EvCM*_). We performed this comparison by calculating the cosine of the angle between the two directions. Specifically, we calculated:

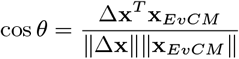

**x** can be a flux or gene vector.

#### D. Non-identifiability of genes

As discussed in the main text, when multiple independent genes can catalyze a reaction, it is unclear how evolution can differentiate among them. To determine if evolution can differentiate among the independent genes, we compared variation at the level of flux to variation at the level of genes. Specifically, we partitioned a simulation across time into separate bins and calculated the change in flux/maximum allowable flux over that bin. Then, for each reaction and gene, we calculated the coefficient of variation of change in flux/maximum allowable flux over the bins. We broadly expected that if there was a one-to-one relationship between genes and reactions, the coefficient of variation should be similar between a gene and its reaction. To examine this, we subtracted the flux-level coefficient of variation from the gene-level coefficient of variation. As we only used simulations where *A* = *I* for this analysis, any gene can easily be associated with a specific reaction. We also only considered one gene for each reaction, as identified by the sign of the fluxes (*i*.*e*., upper genes if the flux is positive and lower genes if the flux is negative).

When we performed this analysis with the toy networks (Figure S4), the results clearly indicate that when multiple genes are present, outcomes at the level of genes are more variable than outcomes at the level of fluxes compared to when a gene and reaction have a one-to-one relationship. Bins of 50,000 attempted mutations were used, so there were 100 bins throughout the simulation. The first bin was not included in analysis to avoid any influence of the initial conditions of the simulation.

When we examined the randomly generated metabolic networks (Figure S8C), estimation of the coefficient of variation was more challenging, and some differences were >100. To avoid these potential outliers influencing the results, we only examined the middle 90% of the data. Outliers were removed across the entire data set, not stratified by the number of other genes for a reaction. Bins of 500,000 attempted mutations were used, so there were 50 bins throughout the simulation. The first bin was not included in analysis to avoid any influence of the initial conditions of the simulation.

---

**Fig. S1:**
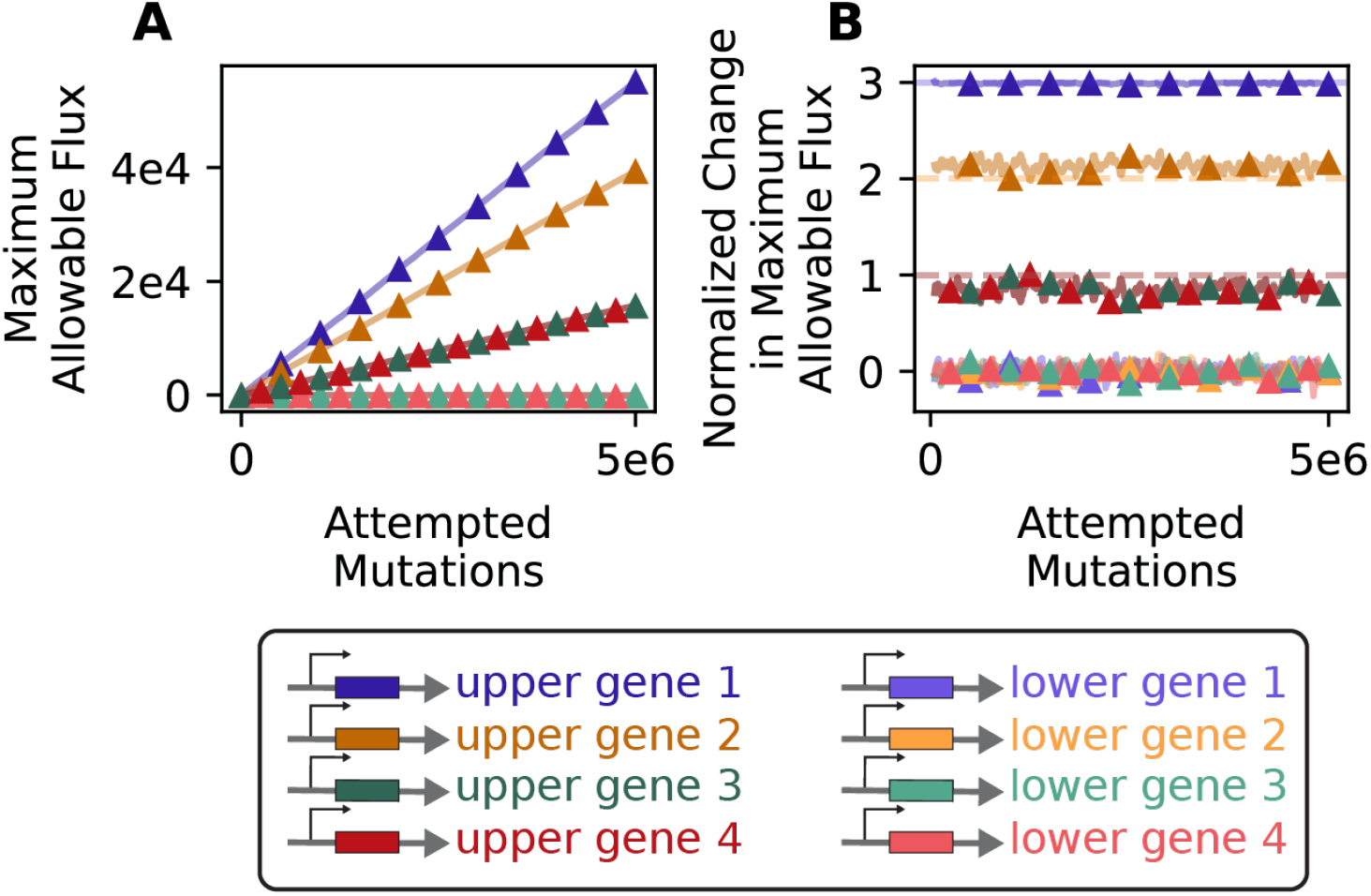
*(A)* The maximum allowable fluxes for the genes of the toy network for the replicate shown in Figure 2. As discussed in Section 3A, upper and lower bounds on reaction fluxes are set by independent genes. *(B)* The normalized changes in maximum allowable flux of each gene is relatively constant, indicating the metabolic networks evolves along a constant direction at the level of genes.

**Fig. S2:**
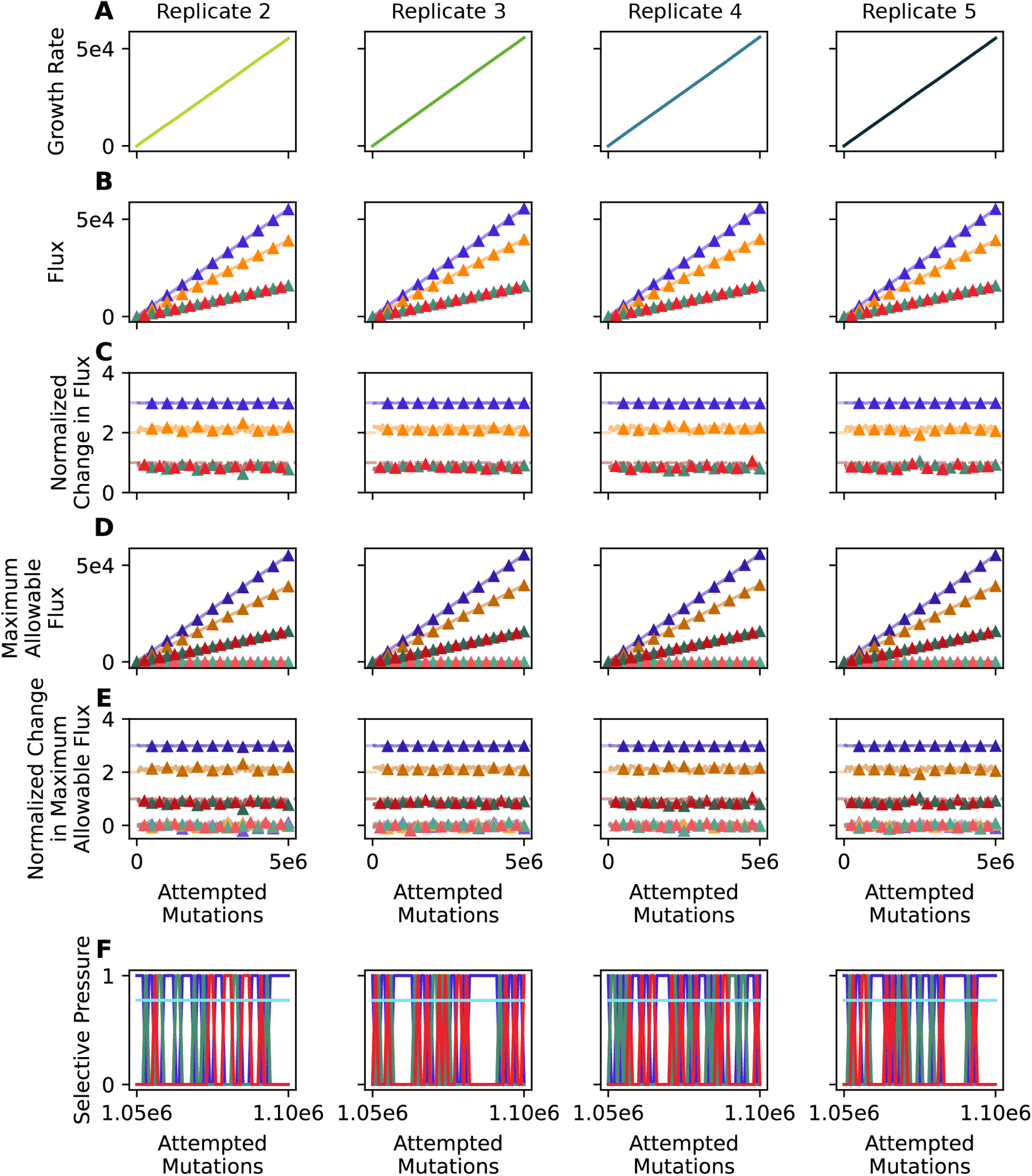
Replicate simulations of the toy network were performed, as discussed in the main text. The results for replicates not in Figure 2 and Figure S1 are shown here. The color of each plot matches the colors in Figure 2 and Figure S1. The growth rate *(A)*, flux *(B)*, normalized change in flux *(C)*, the maximum allowable flux *(D)*, and normalized change in maximum allowable flux are shown for the entire simulation. *(E)* The selective pressure on the individual genes and EvCM are shown for the same period of the simulation as in Figure 1E. As discussed in the main text, the selective pressure on the individual genes varies across time. However, comparison across replicates indicates that this variation is also random; the selective pressure at a given time varies across replicates. This is not true for the collective mode; its selective pressure is constant across time and replicates.

**Fig. S3:**
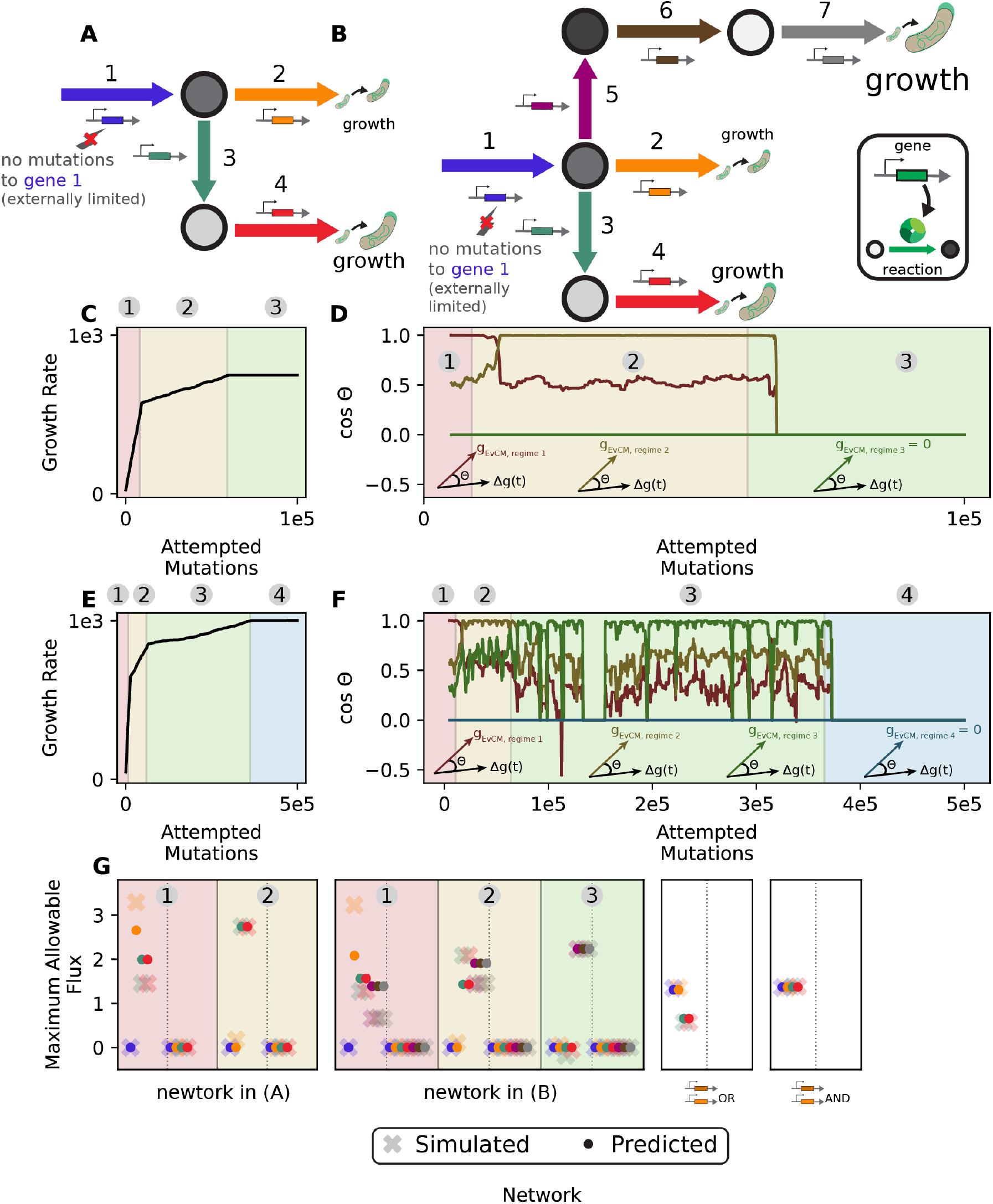
Additional modifications of the toy metabolic network (Section 4B) show how multiple (*>*2) regimes arise. *(A)* The toy network was modified in the same way as Figure 3 (***a***_***max***_ ≤ ∞), except the reaction 2 pathway leads to less growth than the reactions 3/4 pathway. Reactions cannot run in reverse in this network. *(B)* The metabolic network in (A) was further modified by adding a third branch (reactions 5, 6, and 7). This third branch generates more growth than either of the previous two branches. Reactions also cannot run in reverse in this network. *(C)* The growth rate of the metabolic network in (A) throughout an evolutionary simulation. Three evolutionary regimes are present. *(D)* The cosine similarity between direction of change and the EvCM for each regime. The EvCM for the third regime is 0, so the cosine similarity is 0. As each regime ends and the next starts, the direction of evolution becomes less aligned with the EvCM of the earlier regime. Again, three regimes can be seen, each with their own direction of evolution. *(E)* The growth rate of the metabolic network in (B) throughout an evolutionary simulation. FOur evolutionary regimes are present. *(F)* The cosine similarity between direction of change and the EvCM for each regime. The EvCM for the fourth regime is 0, so the cosine similarity is 0. As each regime ends and the next starts, the direction of evolution becomes less aligned with the EvCM of the earlier regime. Again, four regimes can be seen, each with their own direction of evolution. *(G)* Predictions of the EvCM were made using the constant and maximal selective pressure criteria for the metabolic networks shown in this figure (within each regime) and the remaining networks from Figure 2 (2 independent genes and 2 dependent genes). Predictions are shown for both upper and lower bounds for the networks in (A) and (B) and are divided by the dotted line.

**Fig. S4:**
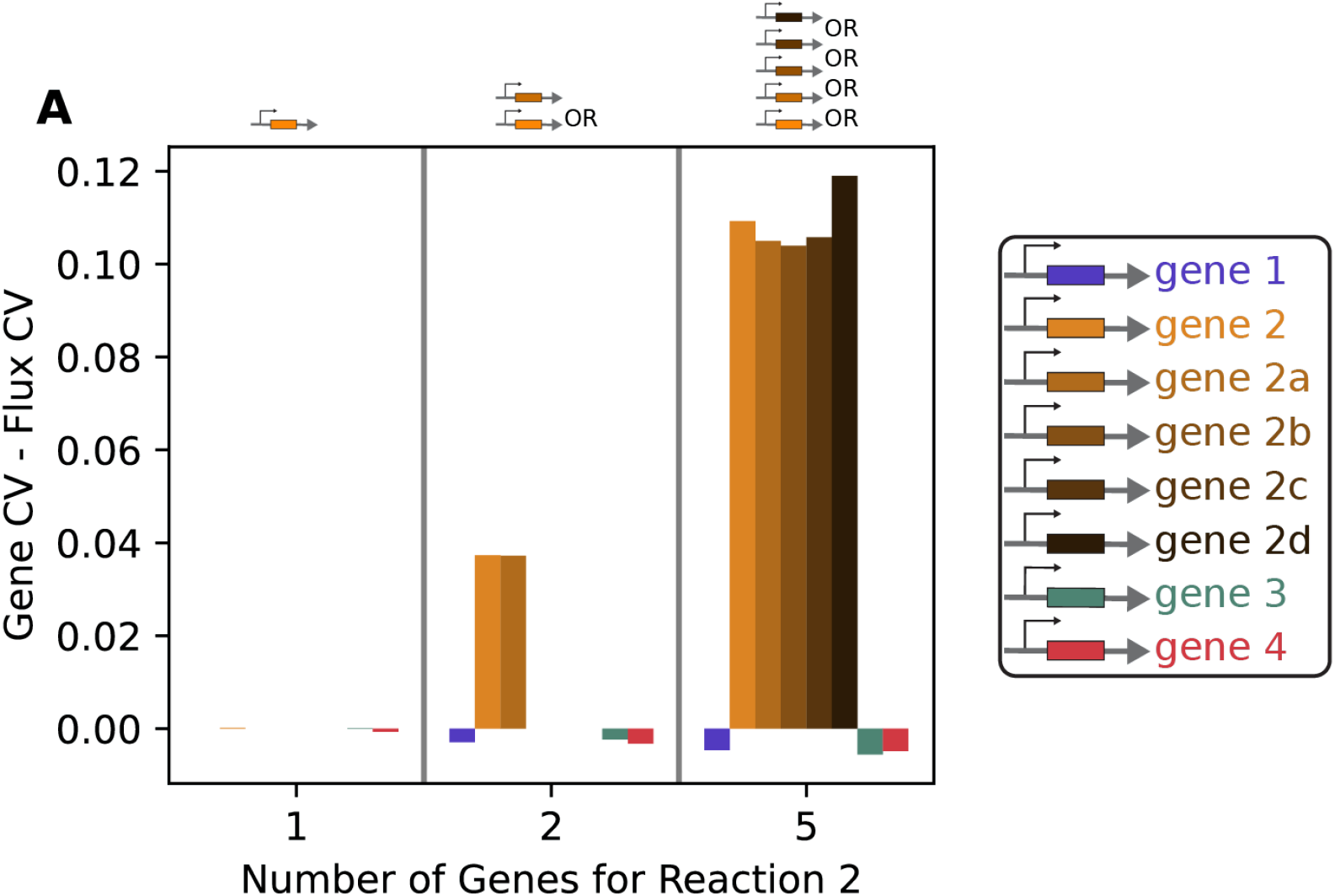
*(A)* The coefficients of variation of the EvCM at the levels of fluxes and genes were calculated for each reaction and gene. Their difference measures the non-identifiability of genes compared to fluxes. As more independent genes are added to reaction 2 (genes 2a, 2b, 2c, and 2d), the difference between gene and flux coefficient of variation increases, indicating that the genes are becoming more non-identifiable. The genes for reactions 1, 3, and 4 have similar gene and flux level coefficients of variation.

**Fig. S5:**
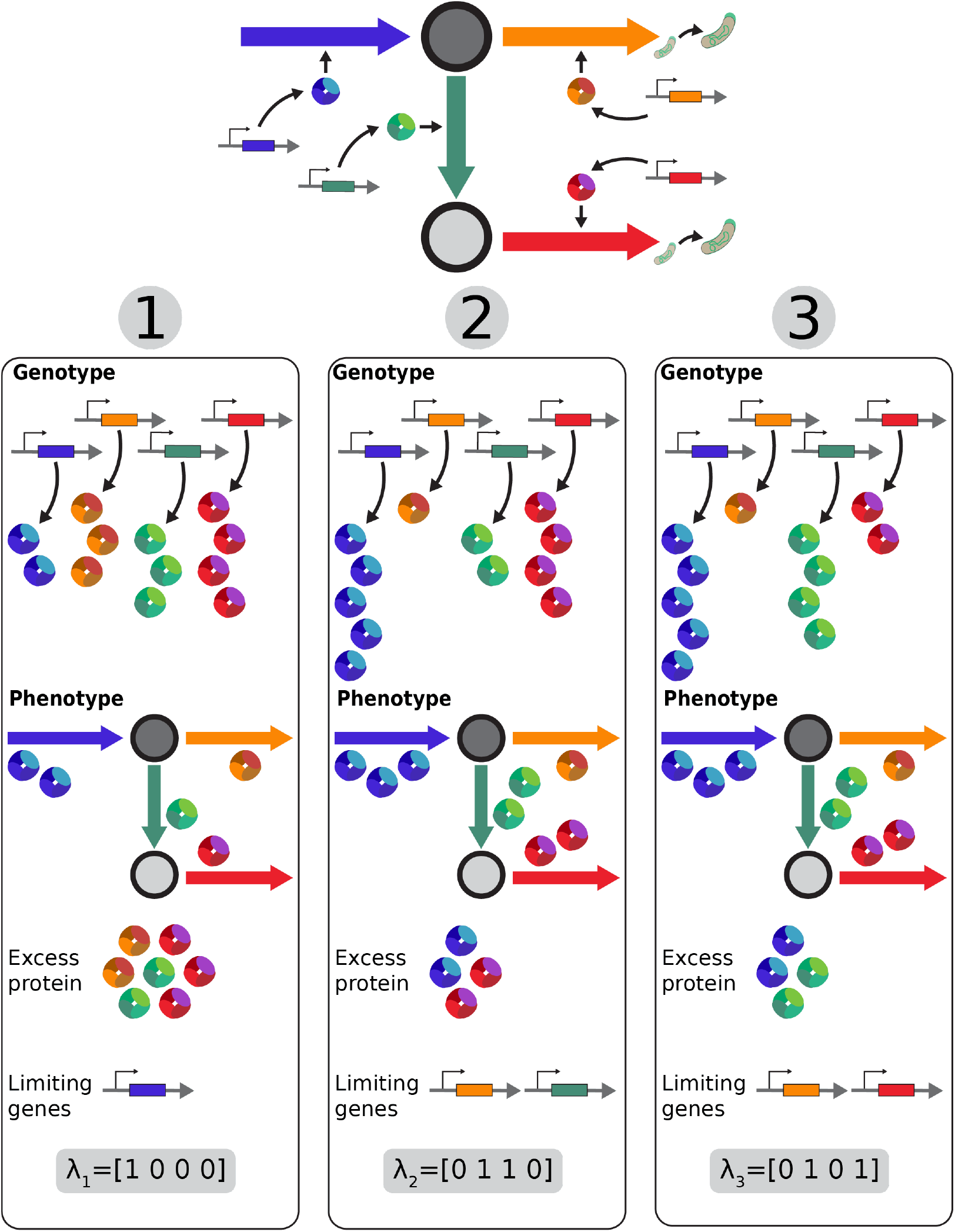
The toy metabolic network has three unique values of ***λ*** as mentioned in the main text, corresponding to three different scenarios. Examples of the genotype and phenotype of these three scenarios are shown schematically in scenarios (1), (2), and (3). In scenario (1), the genotype leads to an approximately equal balance of protein across genes. However, the phenotype must follow conservation of mass, so there is excess protein for reactions 2, 3 and 4. This means that gene 1 is rate-limiting for the network, and so ***λ*** = [1, 0, 0, 0]. Alternatively, in scenario (2), the genotype leads to a large amount of protein for reactions 1 and 4. At the level of phenotype, there is then excess protein for reactions 1 and 4. So, genes 2 and 3 are rate-limiting and ***λ*** = [0, 1, 1, 0]. Scenario (3) is similar to scenario (2), except now there is excess protein for reaction 3 and reaction 4 is rate-limiting. Then, ***λ*** = [0, 1, 0, 1].

**Fig. S6:**
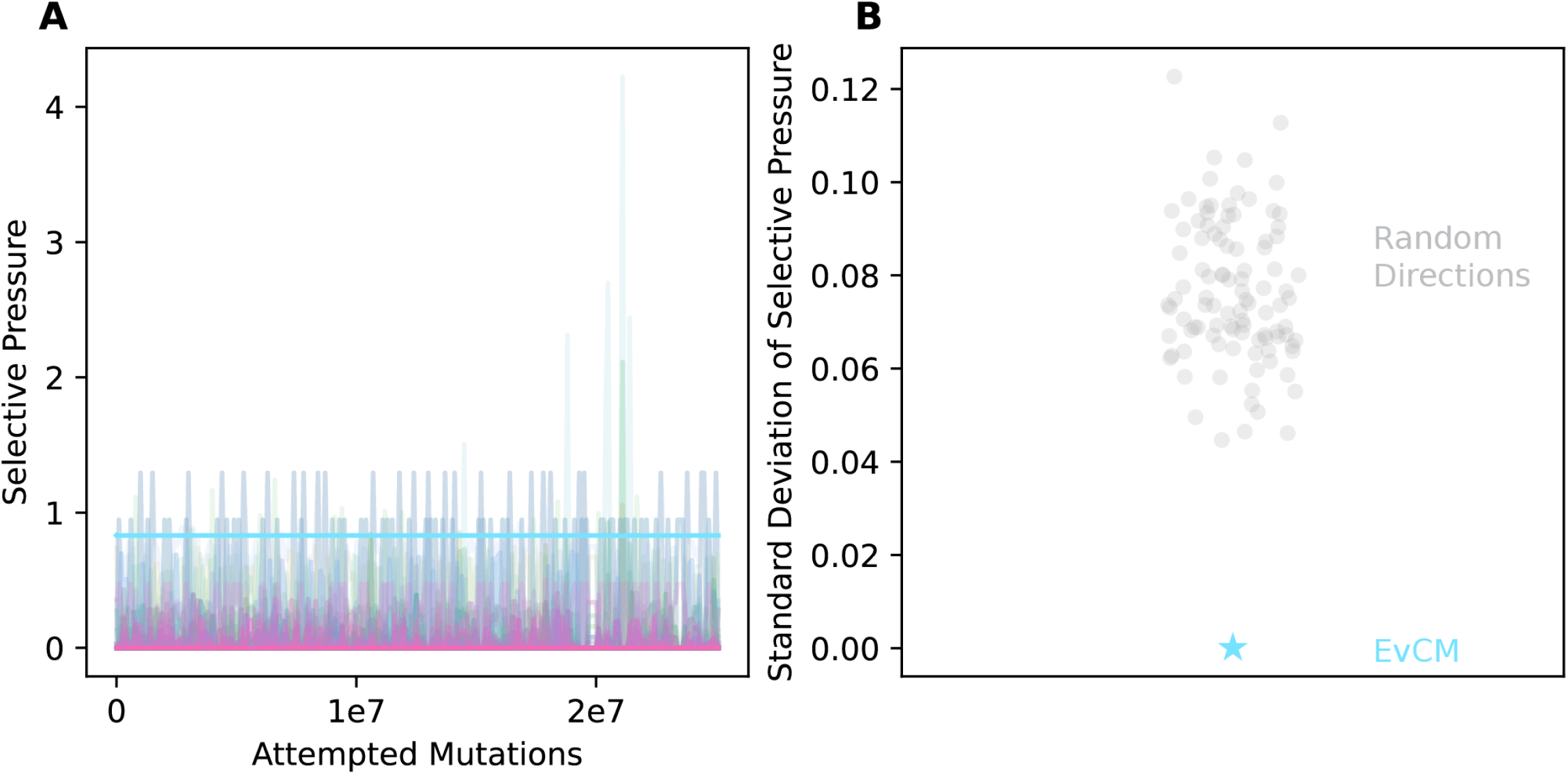
The EvCMs of the random metabolic networks have constant selective pressure, as discussed in the main text. Here, the selective pressure of the EvCM for the random metabolic network shown in Figure 4 is characterized. *(A)* The selective pressure on the EvCM (light blue) and the individual genes (faint lines) throughout the evolutionary simulation. The selective pressure on the EvCM is constant, while the selective pressure on the individual genes varies across time. *(B)* The standard deviation of the selective pressure on the EvCM and random directions. The EvCM has constant selective pressure compared to random directions.

**Fig. S7:**
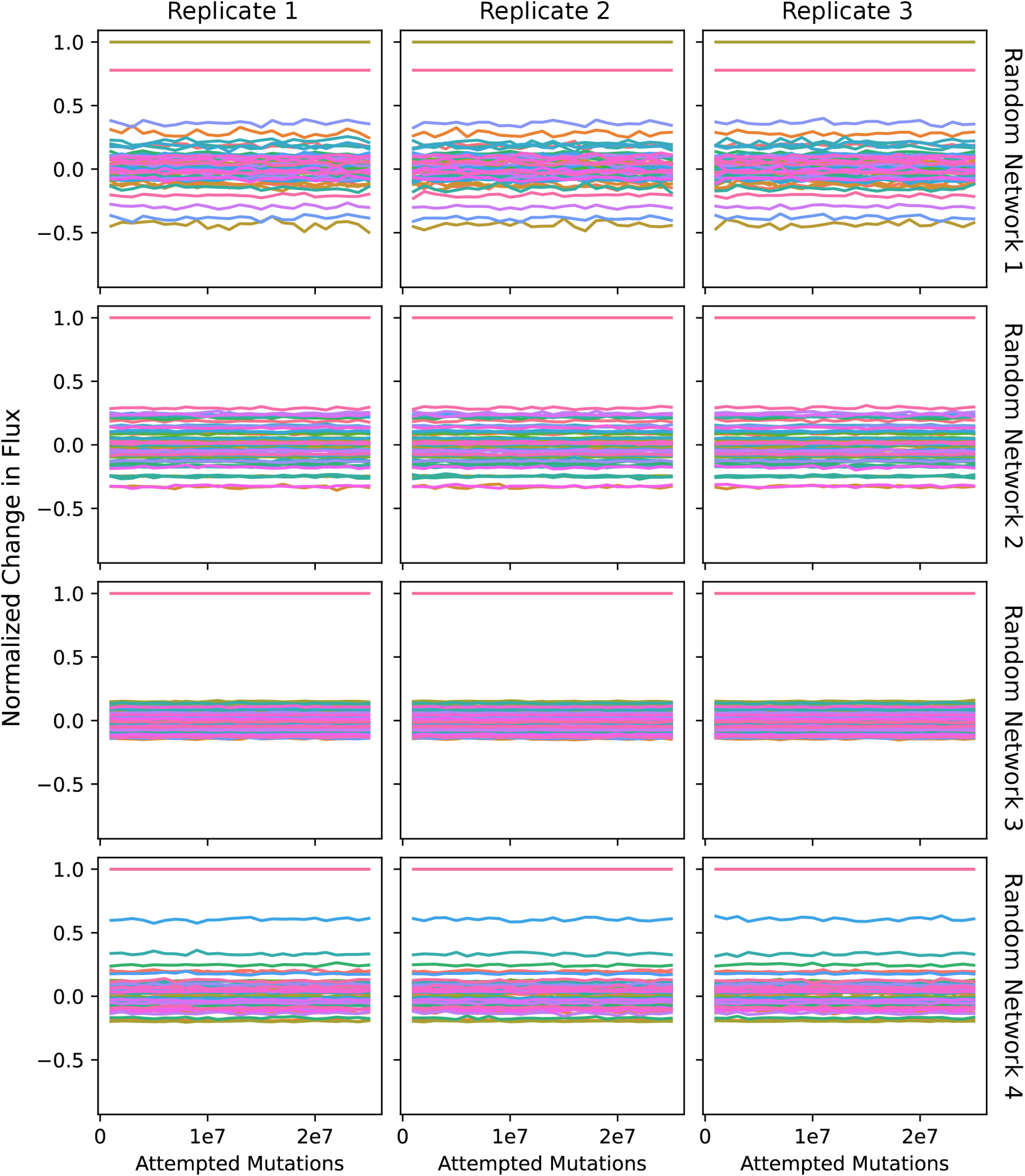
Replicate simulations of random metabolic networks were performed. Each plot shows the normalized change in flux throughout an evolutionary simulation. Each row is an individual random metabolic network, and each column is a replicate of the simulation. The directions of evolution are consistent across time and replicates.

**Fig. S8:**
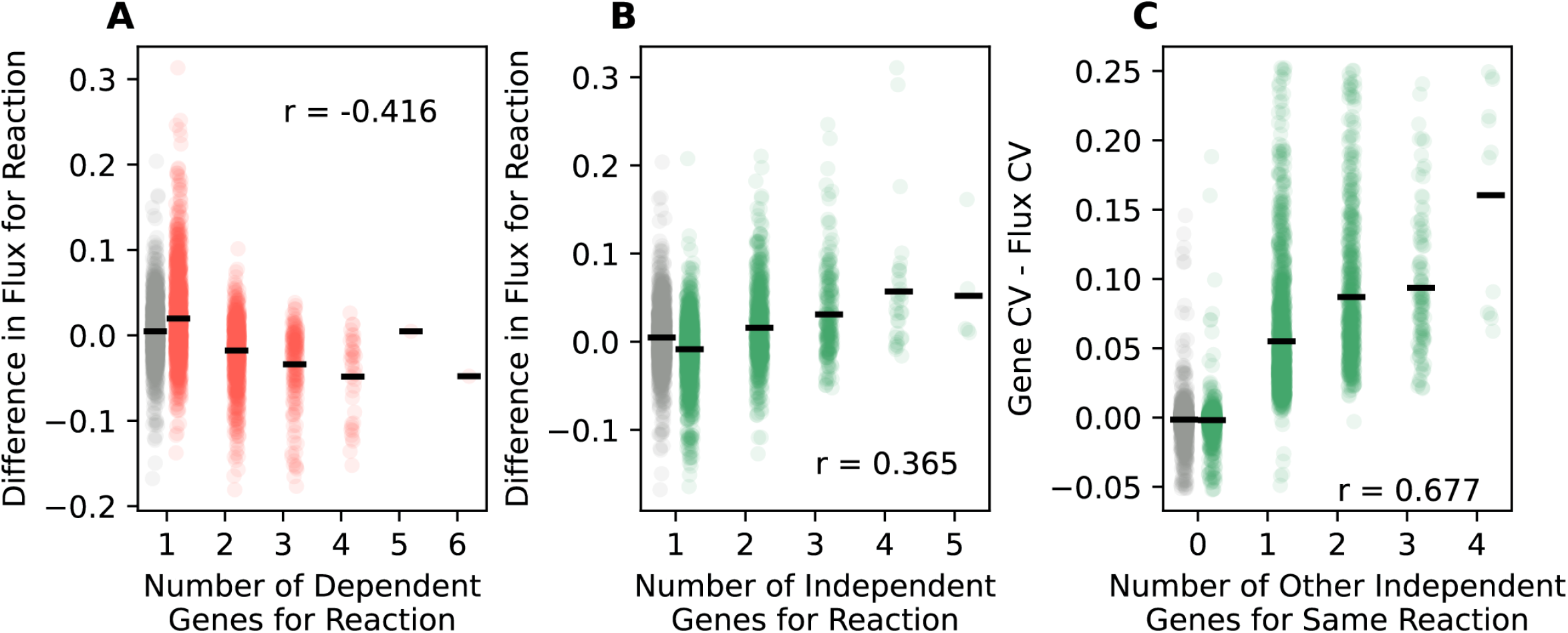
The simulations of the random metabolic networks were analyzed in a similar manner to the toy network (Figures 3E and 3F and Figure S4) to understand what shapes the EvCM and the identifiability of genes. Simulations with where reactions were controlled by a random number of multiple dependent (A) or independent (B, C) genes were performed. For comparison, the data in grey in each plot represents simulation results where the entire metabolic network had one gene per reaction. In all plots, black lines represent means of associated data. In (A) and (B), the EvCM of simulation was compared to the prediction if only one gene coded for each reaction (Section 7C). In (C), analysis was carried out as in Figure S4 (Section 10D). *(A)* The flux in the EvCM allocated to a reaction with multiple dependent genes decreases as the number of dependent genes increases. *(B)* The flux in the EvCM allocated to a reaction with multiple independent genes increases as the number of dependent genes increases. The results of (A) and (B) suggest the EvCM favors more evolvable directions. *(C)* The coefficient of variation of genes compared to the coefficient of fluxes partitioned by the number of other independent genes for the same reaction. The difference increases with the number of other independent genes, suggesting the genes become more non-identifiable.

**Fig. S9:**
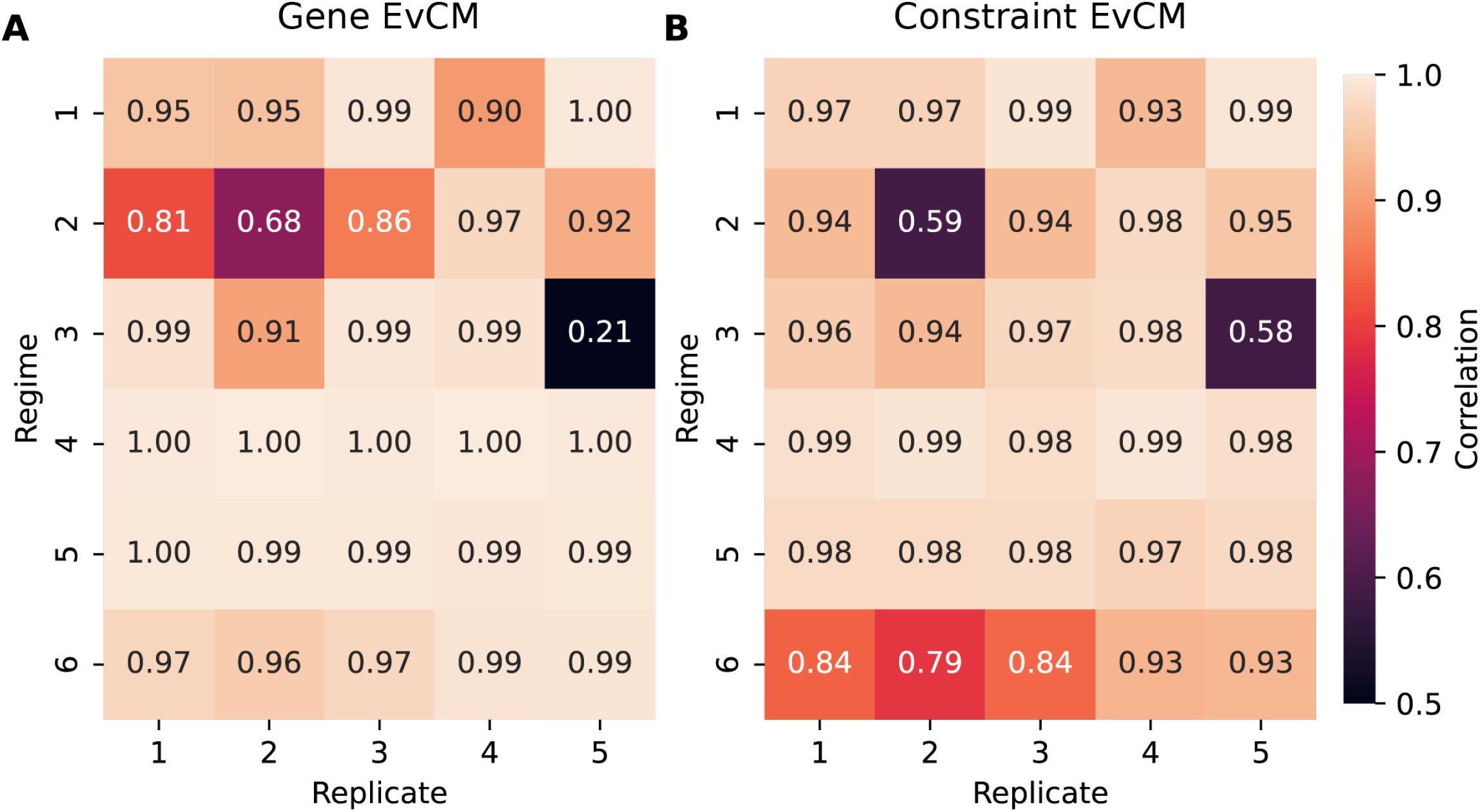
Predictions for the *E. coli* EvCMs were made for each regime and replicate using the values of ***λ*** from simulation (Section 6C) *(A)* Prediction at the level of genes (**z**_*g*_). Predictions at the level of constraints (**z**_*c*_).

**Fig. S10:**
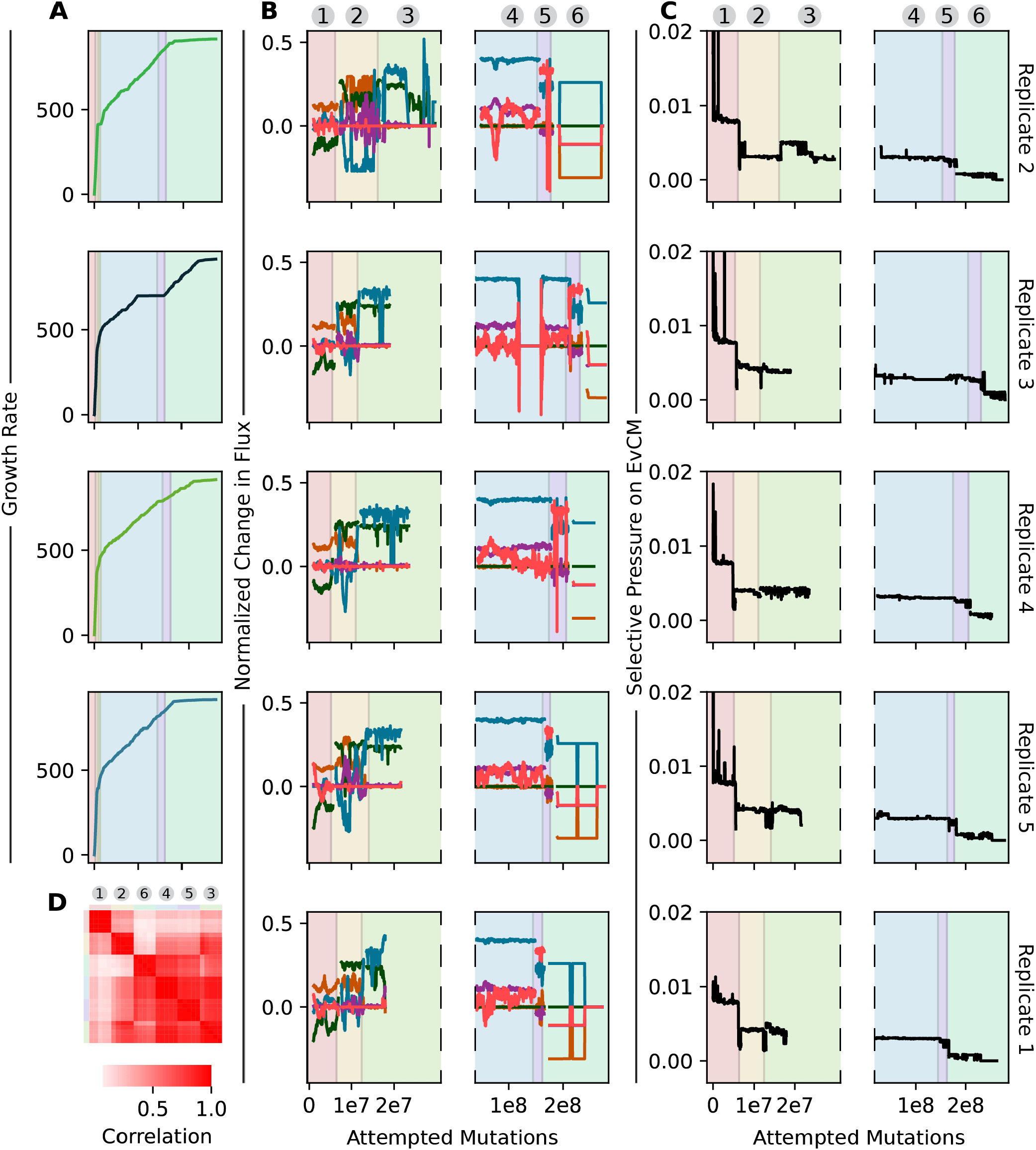
Replicate simulations were performed of the *E. coli* central metabolism. This figure shows results for each individual replicate. In all plots, the regimes are indicated by the background color. *(A)* The growth rates throughout the simulation for each replicate. The growth rate for Replicate 1 is shown in Figure 5B. *(B)* The normalized change in flux within each regime. Directions of evolution are consistent across time (within a replicate) and across replicates. The plots are split in two due to the difference in time scale of the regimes. *(C)* The selective pressure on the EvCM within each regime, for each replicate. The selective pressure is nearly constant within regimes but varies across regimes. Like in (B), the plots are split in two due to the difference in time scale of the regimes. *(D)* The same analysis as in Figure 5C, but at the level of genes. The gene EvCM within each regime was estimated for each replicate simulation. Hierarchical clustering was performed, and the EvCMs cluster by regime, as indicated by color along border, showing consistency across replicates.

**Fig. S11:**
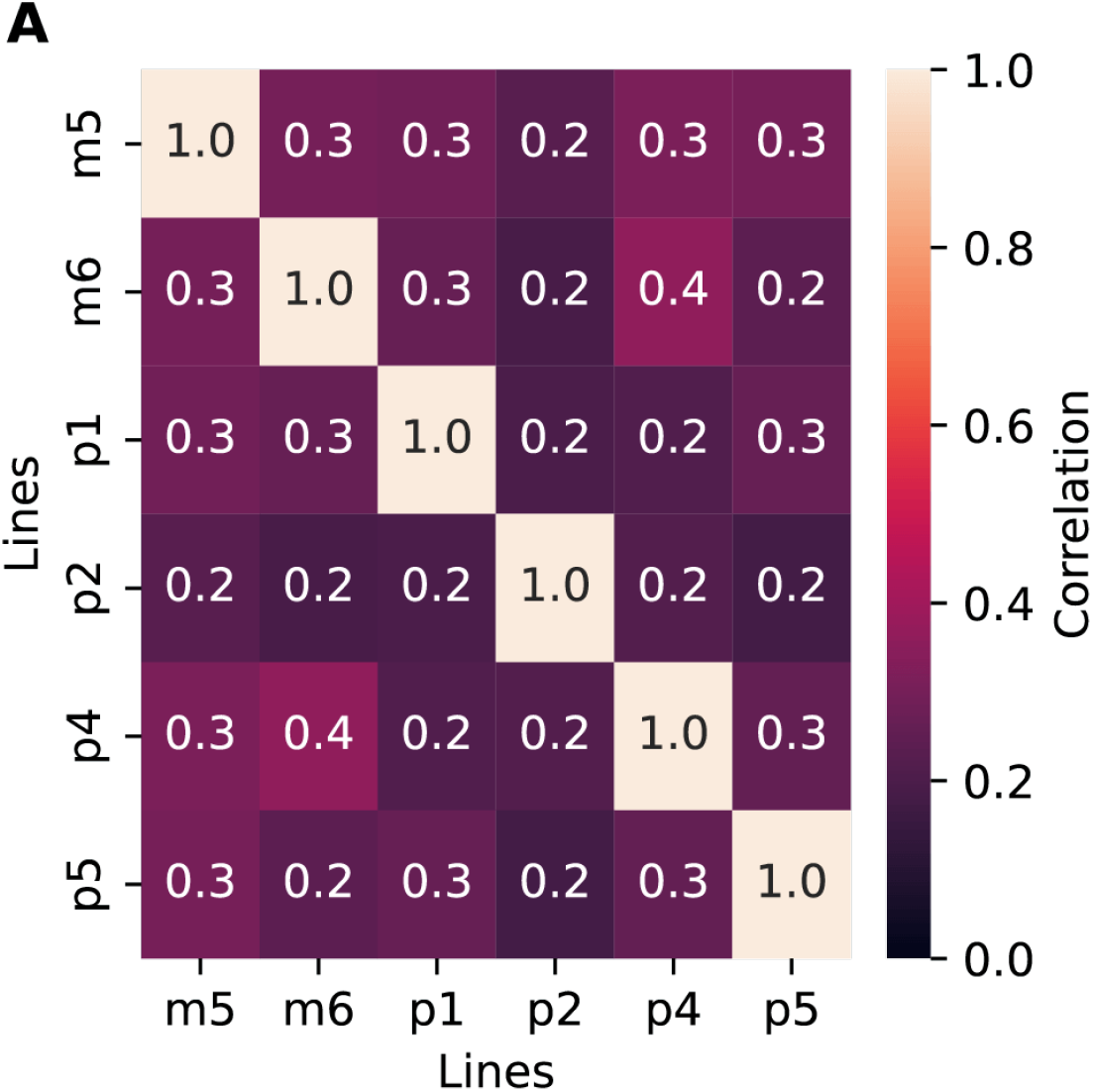
*(A)* The correlations between mutation counts across non-mutator lines. Mutations in each of 3,969 genes were counted across lines, and correlations of these counts were calculated.

**Fig. S12:**
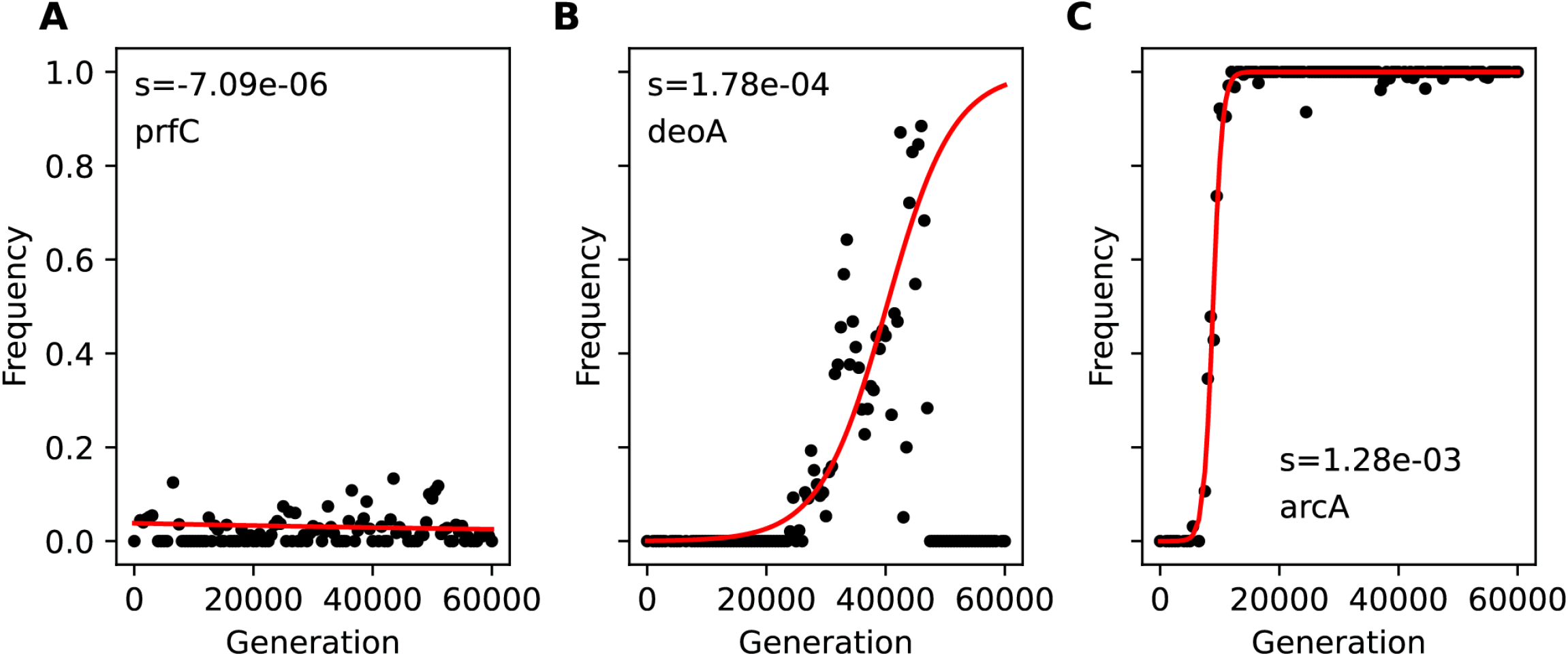
Three examples of logistic fits to mutation frequency trajectories, with the mutated gene indicated. All shown mutations were in the p1 line. *(A)* An example of a nearly neutral mutation. *(B)* An example of how cleaning the data improves the fit. Fits were only performed to the time point at which the maximum frequency occurred, so all data beyond ∼50,000 generations was not used in this example. *(C)* A third example of a fit with a positive selective advantage.

**Fig. S13:**
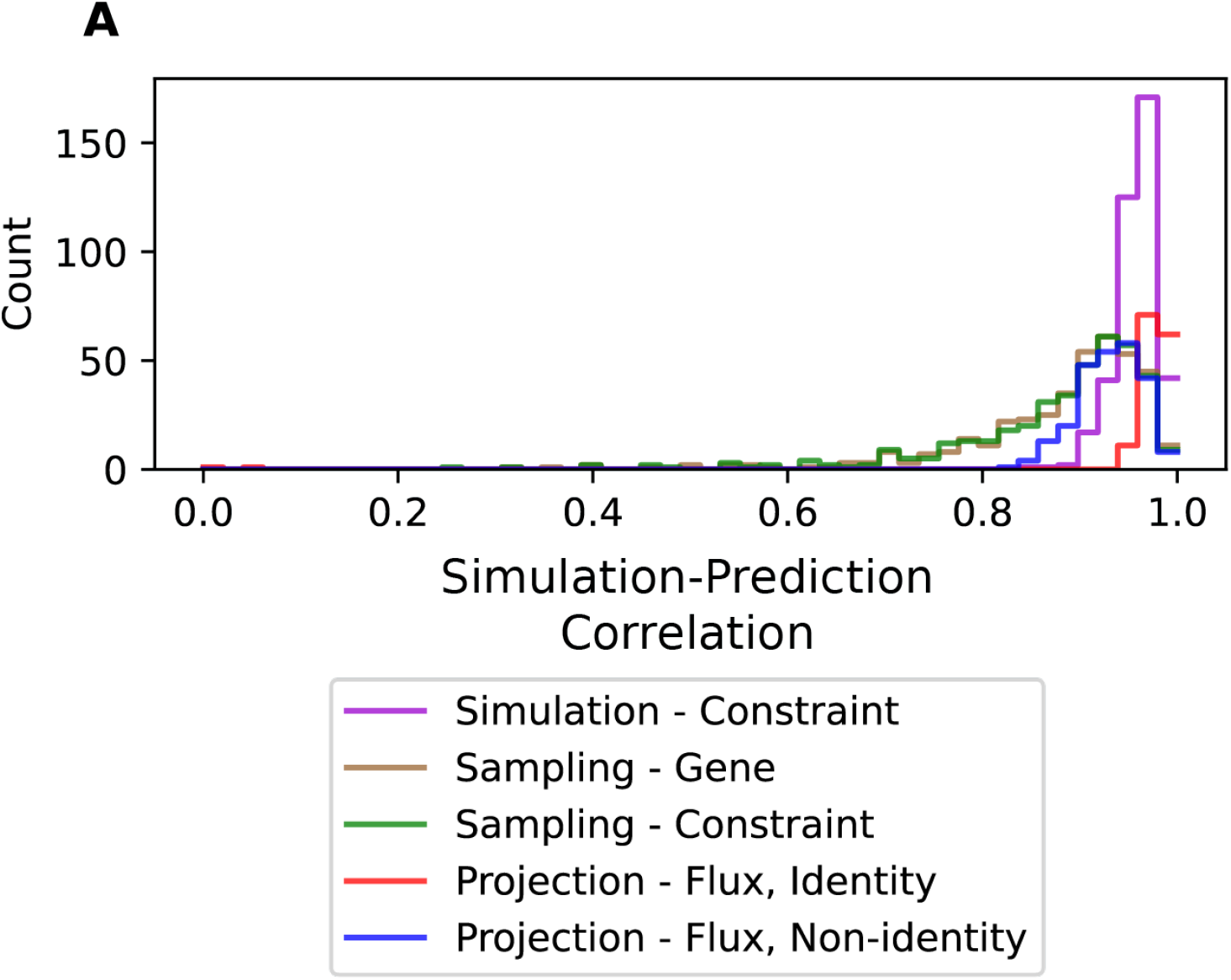
The main text discusses one method to predict the EvCM of random metabolic networks at the level of genes. Other prediction approaches can be used, and predictions can be made as the level of constraints and flux as well (Section 6C). This figure shows the distribution of correlations between simulation and prediction EvCM using various approaches. *(A)* Correlations at the level of constraints (**z**_*c*_) when using values of ***λ*** from simulation. *(B)* Correlations at the level of genes (**z**_*g*_) when using the chain sampling approach. *(C)* Correlations at the level of genes (**z**_*c*_) when using the chain sampling approach. *(D)* Correlations at the level of fluxes (**z**_*f*_) when using Section 6E Equation 15 for matrices with *A* = *I* and *G* = *I. (E)* Correlations at the level of flux (**z**_*f*_) when using Section 6E Equation 15 for matrices with *A* ≠ *I* or *G* ≠ *I*.

**Fig. S14:**
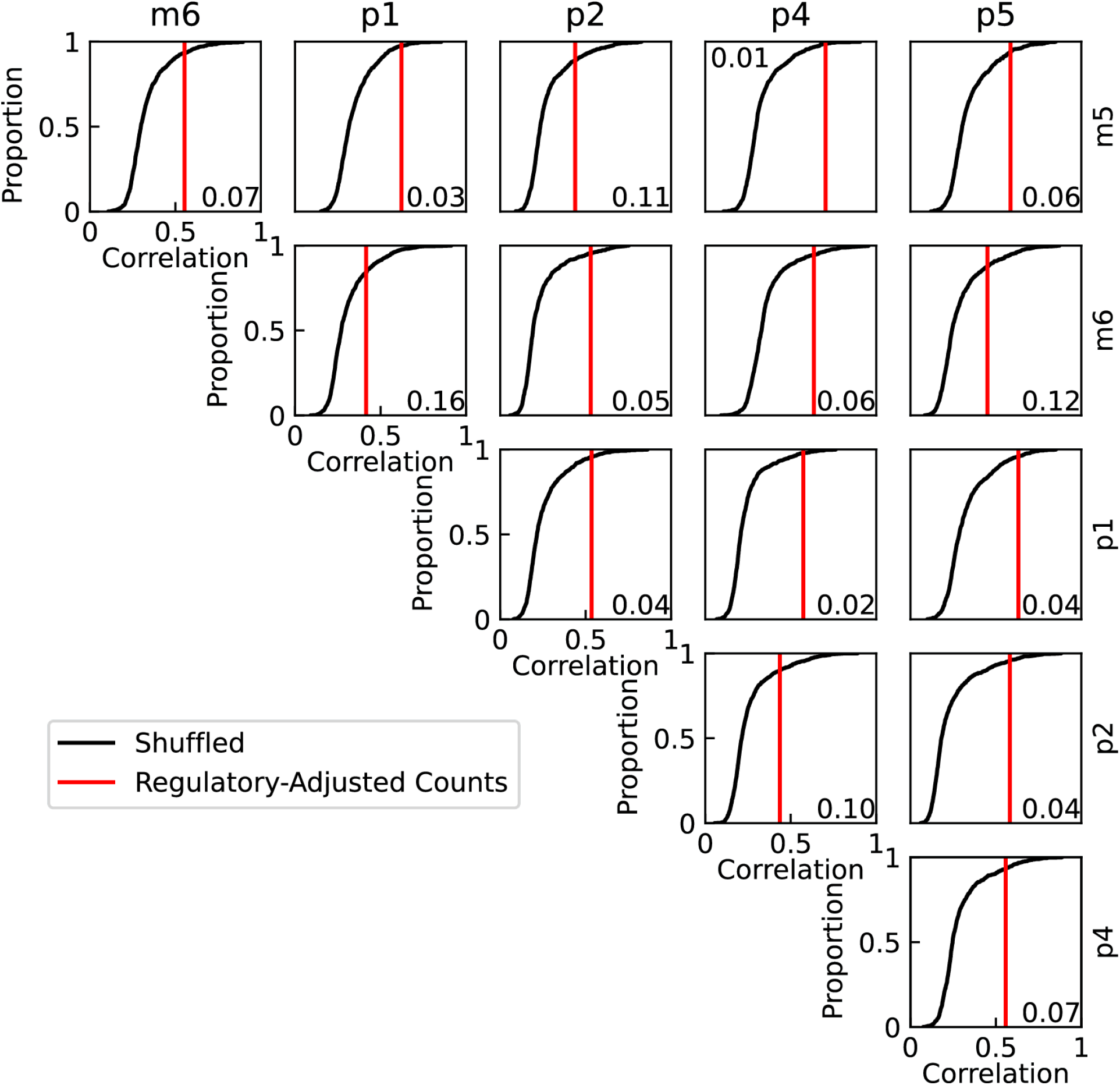
A null distribution was generated (Section 9A) to compare the raw correlations of mutation spectrums across lines (Figure S11) the regulatory-adjusted correlations (Figure 6B). Non-mutator lines are indicated along the top row and right column. The red vertical line indicates the correlation observed (*i*.*e*., the value in Figure 6B), and the black line is the ECDF of 1,000 samples of the null distribution. p-values are indicated within each subplot.

**Table S1.**
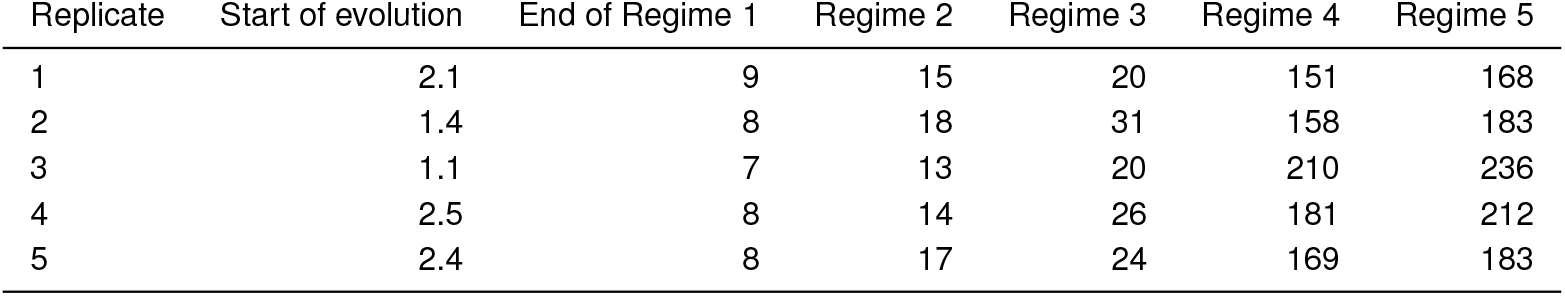
Estimated time step (in millions of time steps) that evolution starts at and each regime ends at in each replicate of the *E. coli* simulations. The simulations were assumed to be in regime 6 after the regime 5 ended.

| Replicate | Start of evolution | End of Regime 1 | Regime 2 | Regime 3 | Regime 4 | Regime 5 |
| --- | --- | --- | --- | --- | --- | --- |
| 1 | 2.1 | 9 | 15 | 20 | 151 | 168 |
| 2 | 1.4 | 8 | 18 | 31 | 158 | 183 |
| 3 | 1.1 | 7 | 13 | 20 | 210 | 236 |
| 4 | 2.5 | 8 | 14 | 26 | 181 | 212 |
| 5 | 2.4 | 8 | 17 | 24 | 169 | 183 |

**Table S2.** Hyperparameters for each group of simulations performed.

| Simulation | Toy network and variants | Random metabolic networks | *E. coli* |
| --- | --- | --- | --- |
| *T* | $5 \times 10^6$ | $> 2 \times 10^7$ | $> 2.7 \times 10^8$ |
| *N* |  | 10,000,000 |  |
| *ss* | 10 | 100 | 10,000 |
| *ms* | 0.1 | 0.1 | 0.01 |
| *em* |  | 1 |  |

**Table S3.** Parameters used to generate random metabolic networks.

| Number of simulations | $m$ | $r$ | $c$ | $h$ | $n_{import}$ | $n_{export}$ | $z_{max}$ | $\nu_{max}$ | $n_{bio}$ | $p$ |
| --- | --- | --- | --- | --- | --- | --- | --- | --- | --- | --- |
| 50 | 25 | 50 | 75 | 75 | 5 | 5 | 5 | 2 | 8 | 1 |
| 50 | 25 | 50 | 49 | 75 | 5 | 5 | 5 | 2 | 8 | 1.5 |
| 50 | 25 | 50 | 49 | 49 | 5 | 5 | 5 | 2 | 8 | 1 |
| 50 | 25 | 50 | 60 | 80 | 5 | 5 | 5 | 2 | 8 | 1.5 |
| 50 | 50 | 100 | 99 | 99 | 10 | 10 | 5 | 2 | 12 | 1 |
| 50 | 50 | 100 | 120 | 140 | 10 | 10 | 5 | 2 | 12 | 1.5 |
| 50 | 100 | 200 | 199 | 199 | 20 | 20 | 5 | 16 | 8 | 1 |
| 50 | 100 | 200 | 230 | 250 | 20 | 20 | 5 | 16 | 8 | 1.5 |

